# Three-dimensional reconstructions of mechanosensory end organs suggest a unifying mechanism underlying dynamic, light touch

**DOI:** 10.1101/2023.03.17.533188

**Authors:** Annie Handler, Qiyu Zhang, Song Pang, Tri M. Nguyen, Michael Iskols, Michael Nolan-Tamariz, Stuart Cattel, Rebecca Plumb, Brianna Sanchez, Karyl Ashjian, Aria Shotland, Bartianna Brown, Madiha Kabeer, Josef Turecek, Genelle Rankin, Wangchu Xiang, Elisa C. Pavarino, Nusrat Africawala, Celine Santiago, Wei-Chung Allen Lee, C. Shan Xu, David D. Ginty

**Author notes:** These authors contributed equally (Annie Handler, Qiyu Zhang).

## Abstract

Specialized mechanosensory end organs within mammalian skin—hair follicle-associated lanceolate complexes, Meissner corpuscles, and Pacinian corpuscles—enable our perception of light, dynamic touch^1^. In each of these end organs, fast-conducting mechanically sensitive neurons, called Aβ low-threshold mechanoreceptors (Aβ LTMRs), associate with resident glial cells, known as terminal Schwann cells (TSCs) or lamellar cells, to form complex axon ending structures. Lanceolate-forming and corpuscle-innervating Aβ LTMRs share a low threshold for mechanical activation, a rapidly adapting (RA) response to force indentation, and high sensitivity to dynamic stimuli^1–6^. How mechanical stimuli lead to activation of the requisite mechanotransduction channel Piezo2^7–15^ and Aβ RA-LTMR excitation across the morphologically dissimilar mechanosensory end organ structures is not understood. Here, we report the precise subcellular distribution of Piezo2 and high-resolution, isotropic 3D reconstructions of all three end organs formed by Aβ RA-LTMRs determined by large volume enhanced Focused Ion Beam Scanning Electron Microscopy (FIB-SEM) imaging. We found that within each end organ, Piezo2 is enriched along the sensory axon membrane and is minimally or not expressed in TSCs and lamellar cells. We also observed a large number of small cytoplasmic protrusions enriched along the Aβ RA-LTMR axon terminals associated with hair follicles, Meissner corpuscles, and Pacinian corpuscles. These axon protrusions reside within close proximity to axonal Piezo2, occasionally contain the channel, and often form adherens junctions with nearby non-neuronal cells. Our findings support a unified model for Aβ RA-LTMR activation in which axon protrusions anchor Aβ RA-LTMR axon terminals to specialized end organ cells, enabling mechanical stimuli to stretch the axon in hundreds to thousands of sites across an individual end organ and leading to activation of proximal Piezo2 channels and excitation of the neuron.

## MAIN TEXT

Our perception of light touch is initiated by activation of mechanically sensitive low-threshold mechanoreceptors (LTMRs) whose cutaneous endings associate with non-myelinating Schwann cells and other cell types to form complex terminal structures called mechanosensory end organs. The structural diversity of LTMR end organs has long been thought to underlie their unique tuning properties^16^. Fast conducting (Aβ) LTMRs can be subdivided by their rate of adaptation to step indentation and vibration tuning properties. The rapidly adapting (RA) LTMRs (Aβ RA-LTMRs) fire only at the onset and/or offset of step indentations and are frequency-tuned, rendering them uniquely suited to detect dynamic tactile features^2, 16, 17^ (Extended Data Fig. 1A-C). In contrast, slowly adapting (SA) LTMRs (Aβ SA-LTMRs) fire throughout the sustained stimulus period of step indentations and exhibit minimal frequency tuning^1, 18, 19^. While the terminal structure of the Merkel cell-neurite complex formed by Aβ SA-LTMRs is relatively simple and the mechanism by which force activates the mechanosensitive ion channel, Piezo2, within this complex is well studied^20–24^, the complex terminal structures formed by the Aβ RA-LTMRs—including the hair follicle lanceolate complex, Meissner corpuscle, and Pacinian corpuscle—and the subcellular localization of Piezo2 within these structures remains unclear. Despite the overall distinct morphologies of the Aβ RA-LTMRs innervating hair follicles, Meissner corpuscles, and Pacinian corpuscles (Fig. 1A), their shared physiological responses to vibrotactile stimuli (Extended Data Fig. 1A-C) suggest a conserved mechanotransduction mechanism underlying light, dynamic touch. Here, we sought to complement an analysis of the subcellular distribution of the requisite mechanically sensitive ion channel Piezo2 with full-scale 3D EM reconstructions of the three Aβ RA-LTMR end organs to generate a general model of mechanotransduction within these structures.

**Figure 1.**
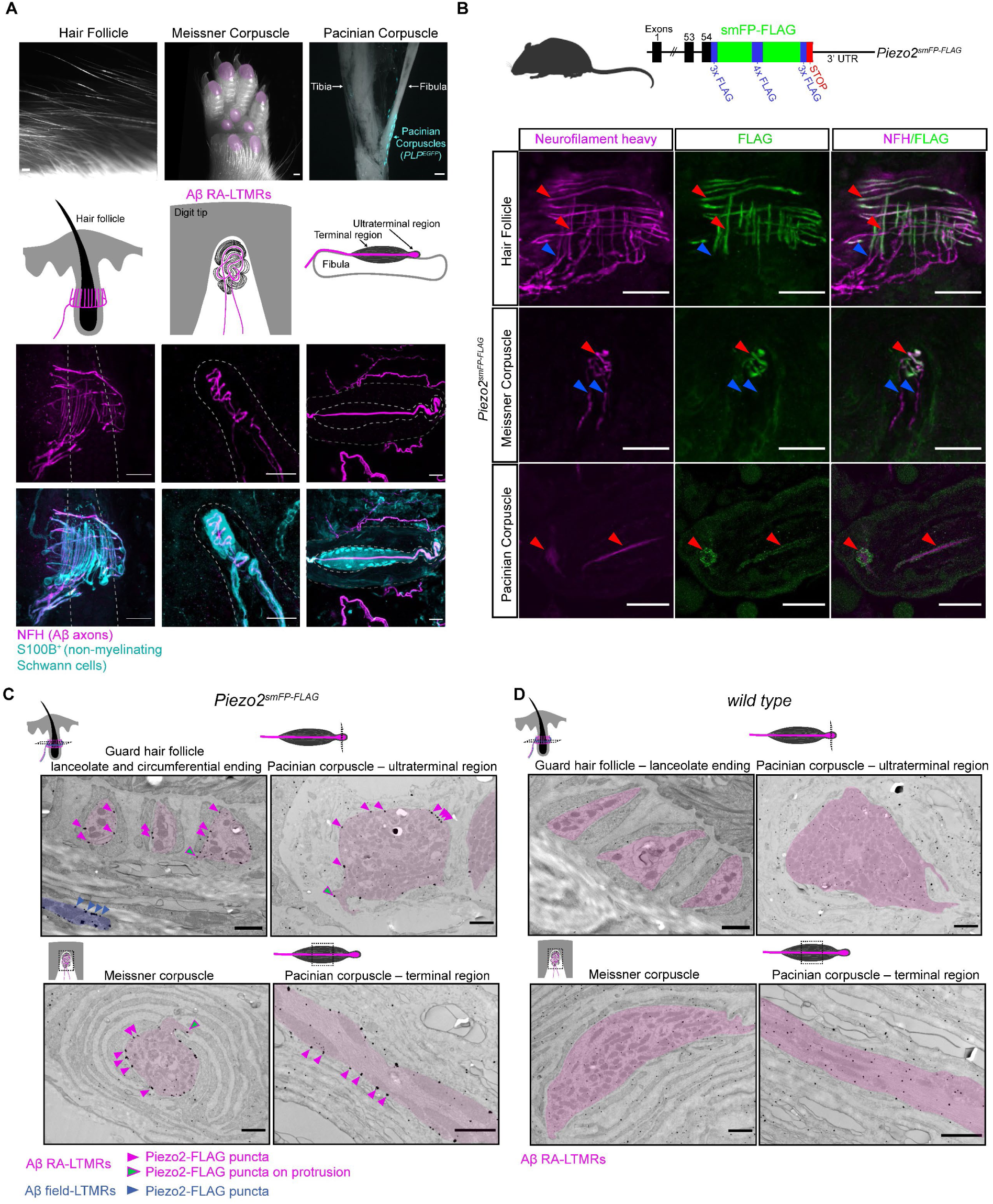
Piezo2 is restricted to axonal endings across morphologically dissimilar Aβ RA-LTMRs. (**A**) Image of the three main skin/tissue regions that contain the end organs formed by Aβ RA-LTMRs, including hairy skin that contains lanceolate endings around hair follicles, glabrous skin that contains Meissner corpuscles within the digit tips and pedal pads (regions indicated in pink), and the periosteum around the fibula that contains Pacinian corpuscles (indicated by the PLP^eGFP^ fluorescent label). Diagrams illustrate end organs with the Aβ RA-LTMRs in magenta. Representative confocal images show the Aβ RA-LTMRs and the specialized non-neuronal cells of each end organ labeled by immunostaining with NFH^+^ and S100B^+^, respectively. Dashed lines indicate the outline of the hair follicle (left), dermal papilla (middle), and the boundary of the outer and inner core (right). Scalebar, 20 µm. (**B**) Top, schematic diagram of the *Piezo2^smFP-FLAG^* allele. Bottom, Piezo2-FLAG fusion protein localizes to the terminal ends of Aβ RA-LTMR axons labeled by NFH^+^ innervating the hair follicle, Meissner corpuscle, and Pacinian corpuscle in *Piezo2^smFP-FLAG^* animals. Red arrowheads point to examples of co-localized FLAG and NFH signal. Blue arrowheads point to NFH signal outside the end organ that lacks FLAG signal. This experiment was repeated in two animals with littermate controls performed side-by-side. Scalebar, 25 µm. (**C**) Immuno-electron micrographs from a *Piezo2^smFP-FLAG^* animal stained for FLAG and processed with silver enhancement show enriched expression of the Piezo2-FLAG fusion protein along the axon membranes of the Aβ RA-LTMRs that form lanceolate endings around the hair follicle and innervate the Meissner corpuscle and Pacinian corpuscle within both the terminal and ultraterminal region (Aβ RA-LTMRs pseudo-colored pink). Piezo2-FLAG puncta can be seen along the sensory axon membranes (pink arrowheads) and along small cytoplasmic axon protrusions (green arrowheads). Piezo2-FLAG is expressed along the membranes of Aβ field-LTMRs that form circumferential endings surround the guard hair (blue). Not all puncta are labeled by arrowheads. This experiment was repeated in two animals with littermate controls performed side-by-side. Scalebar, 1 µm. (**D**) Immuno-electron micrographs from a wild-type littermate of the *Piezo2^smFP-FLAG^*animal shown in (C) stained for FLAG and processed with silver enhancement. The Aβ RA-LTMRs of each end organ are pseudo-colored pink. Scalebar, 1 µm.

### Piezo2 localization across three Aβ RA-LTMR end organ structures

Because tactile-evoked responses begin at the site of mechanotransduction, we first localized Piezo2 within the terminal structures innervated by Aβ RA-LTMRs—the guard hair follicle, Meissner corpuscle, and Pacinian corpuscle. We generated a knock-in mouse in which a “spaghetti monster-FLAG” (smFP-FLAG) epitope tag^25^ is fused to the carboxy terminal end of the Piezo2 protein (*Piezo2^smFP-FLAG^* mice; Fig. 1B) to enable specific, high affinity immuno-localization of endogenous Piezo2. The multimerized FLAG epitope inserted into the scaffold of GFP proved beneficial in enhancing the signal-to-noise in detecting Piezo2 in the skin beyond what was possible with a published Piezo2 reporter line^7, 20^. When hairy and glabrous (non-hairy) skin of *Piezo2^smFP-FLAG^* mice were co-stained with anti-FLAG and anti-neurofilament heavy chain (NFH, a reporter for Aβ axons), we observed spatial colocalization of Piezo2-FLAG and NFH in Aβ sensory axons forming lanceolate and circumferential endings around hair follicles in hairy skin, Meissner corpuscles in glabrous skin, and Pacinian corpuscles associated with bone (Fig. 1B and Extended Data Fig. 2A-B). While Piezo2 was localized to the terminal portion of Aβ axons forming the end organ structures, it was noticeably absent from NFH^+^ axon fibers before they entered the end organ structures (Fig. 1B and Extended Data Fig. 2A). Piezo2 was also absent from resident non-neuronal cells, most easily appreciated in the lack of Piezo2-FLAG labeling in the S100^+^ spheres formed by lamellar cells around Meissner afferents and the GFP^+^ spheres formed by PLP^eGFP+^ lamellar cells in Pacinian corpuscles (Extended Data Fig. 2A). Strikingly, we found small Piezo2^+^, spine-like protrusions (<1 µm in length) that emanated from the NFH^+^ axons (Extended Data Fig. 2A) forming the lanceolate endings, Meissner corpuscles, and Pacinian corpuscles, hinting at the presence of ultrastructural features of Aβ RA-LTMRs that may be involved in mechanotransduction.

**Figure 2.**
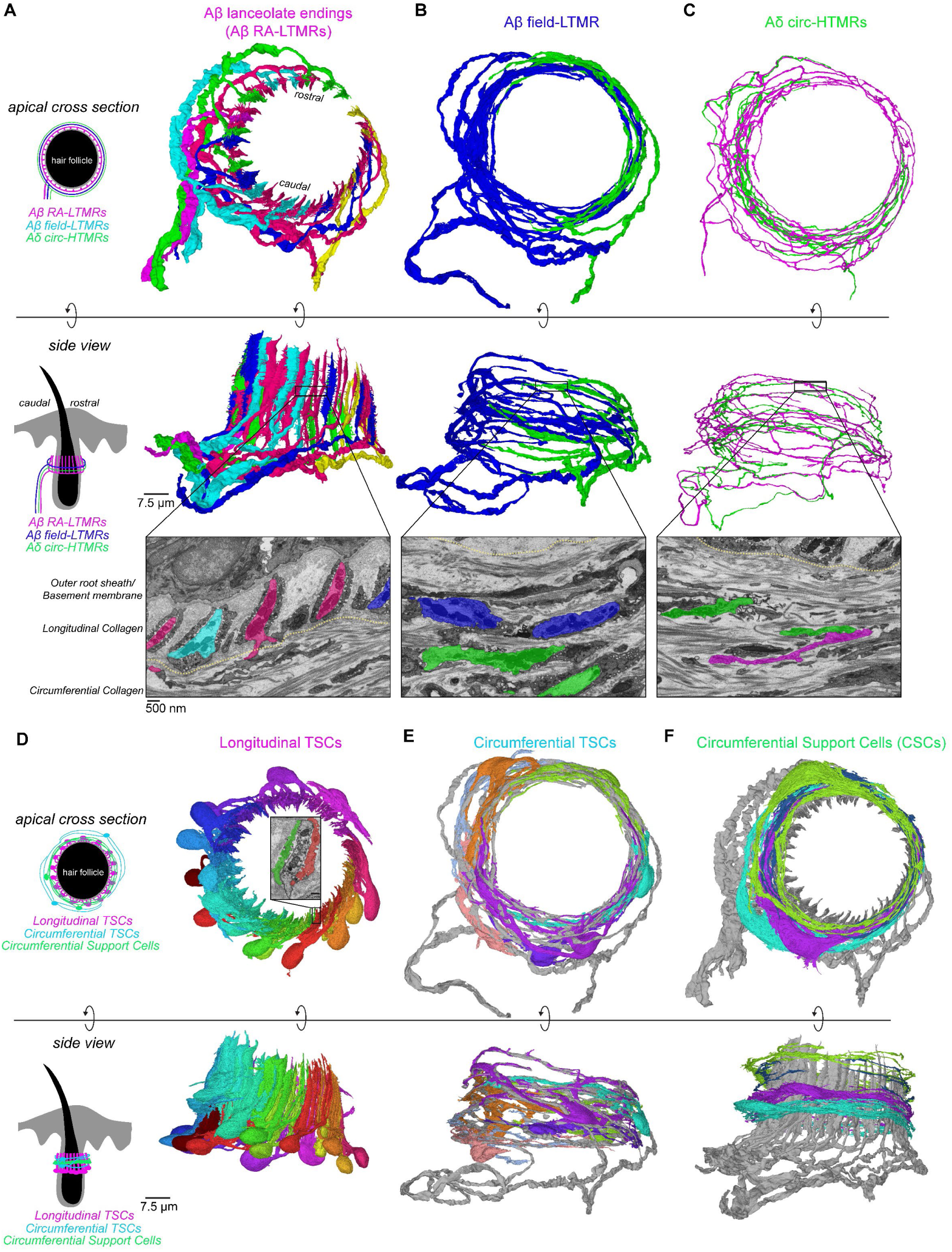
FIB-SEM reconstructions of Aβ and Aδ sensory neuron endings and associated non-neuronal cells of a guard hair follicle. (**A-C**) 3D renderings of sensory neurons innervating a mouse guard hair follicle shown from an apical, cross-section view through the hair follicle (top panels, rostral/caudal axis is indicated) and from a side view of the hair follicle (middle panels). The bottom panels show pseudo-colored images from the FIB-SEM volume with the boundary between the longitudinal and circumferential collagen matrix marked by the yellow, dotted line. (**A**) Six Aβ RA-LTMR axon segments interdigitate within the longitudinal collagen matrix to form 47 lanceolate endings surrounding the guard hair follicle. (**B**) Two Aβ field-LTMRs and (**C**) two Aδ circ-HTMRs innervate the circumferential collagen that lies just beyond the longitudinal collagen network. These Aβ field-LTMRs and Aδ circ-HTMRs branch extensively within the circumferential collagen matrix and form circumferential endings around hair follicle. Additional small diameter axons in the circumferential collagen were seen but their restricted innervation pattern was dissimilar to the anatomical characteristics of the Aδ circ-HTMR. (**D-F**) 3D renderings of non-neuronal cells that associate with the sensory neurons of the guard hair. Top panels show an apical, cross-section view through the hair follicle and bottom panels show a side view of these non-neuronal cells. (**D**) 17 terminal Schwann cells (TSCs) reside within the longitudinal collagen matrix to form mostly tiled, intimate associations with the lanceolate endings. The inset shows an example of two different TSCs associated with the same lanceolate ending. (**E**) 3D renderings of seven circumferential TSCs that reside within the circumferential collagen matrix and associate with the two Aβ field-LTMRs (gray). These seven circumferential TSCs are a significant subset of the total circumferential TSCs present in the volume. (**F**) 3D renderings of four circumferential support cells (CSCs) that also reside within the circumferential collagen matrix. These cells have minimal interaction with the circumferential sensory endings, but frequently contact the axon protrusions of Aβ RA-LTMR lanceolate endings (gray). These four cells are a subset of roughly 17 CSCs within the volume that reside on the same horizontal plane as the lanceolate endings.

Aβ RA-LTMR axons interact with non-myelinating Schwann cells referred to as terminal Schwann cells (TSCs) in hairy skin and lamellar cells in Meissner corpuscles and Pacinian corpuscles. Whether these cells serve as passive modulators of mechanical forces impinging on the sensory axon or play an active role in initiating mechanical responses is an area of active investigation^26–28^. Deleting Piezo2 in both somatosensory neurons and non-neuronal cells leads to complete loss of light touch responses measured in the dorsal root ganglia (DRG), spinal cord, and brainstem, emphasizing the critical role of Piezo2 in light touch^10, 11^. However, definitively answering whether Piezo2 is expressed and functions within end organ non-neuronal cells requires a high-resolution Piezo2 localization approach and physiological analyses of mice lacking *Piezo2* only in non-neuronal cells. Therefore, we next performed immunoelectron microscopy with samples from *Piezo2^smFP-FLAG^* animals to determine the cell-type specific and subcellular localization of Piezo2 within the end organs. We observed Piezo2-FLAG enriched along the sensory axon membranes in all three end organs (Fig. 1C and Extended Data Fig. 3A), consistent with our light microscopy findings. Piezo2-FLAG was localized to Aβ RA-LTMR membranes of hair follicle lanceolate endings, Aβ field-LTMR circumferential endings surrounding hair follicles, and Aβ LTMR terminals in Meissner corpuscles and Pacinian corpuscles (Fig. 1C and Extended Data Fig. 3A). Similar to the Piezo2^+^ spine-like processes observed by light-microscopy, we also observed Piezo2-FLAG localization along small cytoplasmic protrusions that extend beyond the main body of the Aβ LTMR axons (Fig. 1C and Extended Data Fig. 3A), suggesting that these curious finger-like structures—first described over fifty years ago^29–33—may^ participate in mechanical gating of the channel. In contrast, little if any immunostaining signal colocalized to the membranes of TSCs and lamellar cells above the level of background staining observed in non-transgenic controls (Fig. 1D and Extended Data Fig. 3B). Consistent with this observation, electrophysiological recordings in *Dhh-Cre; Piezo2^f/f^* mice showed that deleting Piezo2 in TSCs and lamellar cells, but not in neurons, did not affect mechanosensitivity or the response properties of Aβ sensory neurons (Extended Data Fig. 4A-E). This is in contrast to the near-complete loss of low-threshold responses in the Aβ sensory neurons of *Cdx2-Cre; Piezo2^f/f^* mice, which lack Piezo2 in both somatosensory neurons and non-neuronal cells (Extended Data Fig. 4B). Taken together, our light microscopy, immuno-EM, and genetic manipulations and electrophysiological findings indicate that in all three end organs the Aβ RA-LTMR sensory axons serve as the main site of Piezo2-dependent mechanotransduction, consistent with previously published data^7, 8, 34^. Moreover, the subcellular distribution of Piezo2 and its localization to axon terminals, including the axon protrusions extending from the sensory axons, suggest the possibility of a common mode of neuronal activation across the morphologically distinct Aβ RA-LTMR end organs.

**Figure 3.**
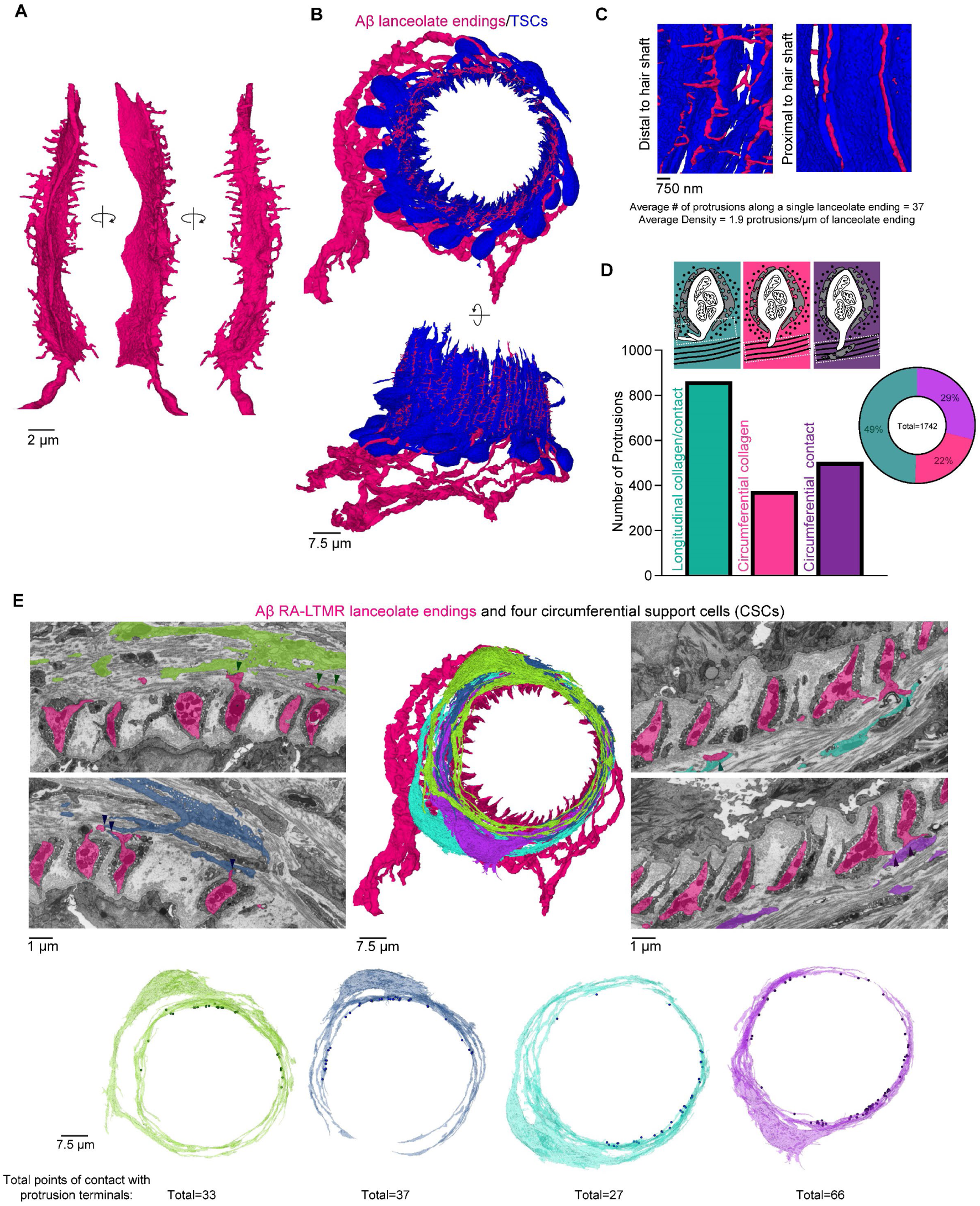
Axon protrusions emerge from Aβ RA-LTMRs, extend through TSC openings, and contact CSCs within the guard hair’s circumferential collagen network. (**A**) A single lanceolate ending with numerous axon protrusions shown from multiple perspectives. (**B**) Apical and side views of the Aβ RA-LTMRs (magenta) and their associated TSCs (blue) reveal extensive axon protrusions that emerge from small gaps in TSC coverage. (**C**) Axon protrusions emerge exclusively at the TSC gaps that occur distal to the hair shaft. Axon protrusions do not emerge from the gaps in TSCs occurring proximal to the hair shaft. The average number of protrusions per lanceolate ending and the density of protrusions along lanceolate endings is indicated. (**D**) Quantification of axon protrusion terminals. Teal: protrusions that stay within the local longitudinal collagen matrix. Magenta: protrusions that extend into the circumferential collagen matrix but do not contact cells in the circumferential collagen network. Purple: protrusions that extend into the circumferential collagen matrix and contact cells in the circumferential collagen network. (**E**) The axon protrusions of Aβ RA-LTMRs that extend into the circumferential collagen network frequently contact the four reconstructed CSCs (arrow heads). Points where each CSC contacts an axon protrusion are marked as dots on the 3D rendering of the cell. The total number of contacts for each cell is noted.

**Figure 4.**
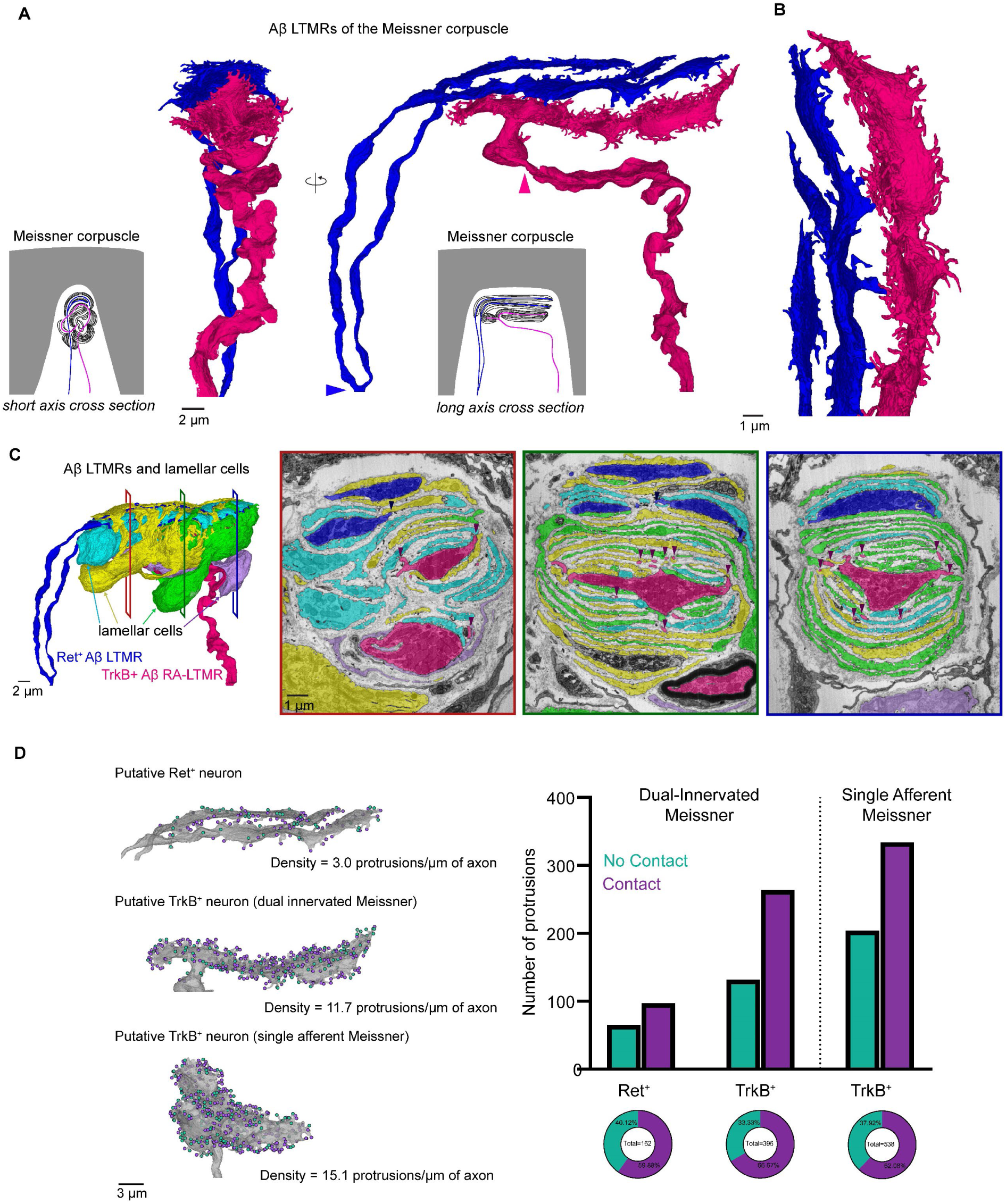
FIB-SEM reconstructions of two Meissner corpuscles reveal the density of axon protrusions and their contacts with corpuscle lamellar cells to be structural correlates of Aβ LTMR tactile sensitivity. (**A**) 3D renderings of two Aβ LTMRs that innervate a single Meissner corpuscle. The arrowheads indicate the position where myelination ends for each axon. (**B**) Higher-magnification rendering of the two Aβ LTMRs. (**C**) Left, 3D renders of the two Aβ LTMRs and the four lamellar cells that form the Meissner corpuscle. Right, pseudo-colored, raw images from the FIB-SEM volume at different depths across the corpuscle. Arrowheads show points where axon protrusions contact lamellar cells. The superficial, blue axon is classified as the putative Ret^+^ Aβ LTMR and the central, magenta axon is classified as the putative TrkB^+^ Aβ RA-LTMR based on the central axon’s more elaborate associations with lamellar cell wrappings throughout the corpuscle. (**D**) Characterization of protrusion terminals of the putative Ret^+^ and TrkB^+^ neurons of the dual-innervated Meissner corpuscle and of the putative TrkB^+^ neuron of a single-innervated Meissner corpuscle from a second FIB-SEM volume. Left, 3D renderings of the two axons from the dual-innervated corpuscle and of the axon from the single-innervated corpuscle. Teal dots indicate protrusions that terminate in the collagen. Purple dots indicate protrusions that contact lamellar cells. The density of protrusions is noted. Right, quantification of protrusion terminals along the sensory axons of the two Meissner corpuscles.

### 3D architecture of three Aβ RA-LTMR end organs

Key insights into mechanotransduction in the cochlea were revealed by classical 3D scanning electron microscopy analysis of cochlear hair cells. This 3D analysis led to the discovery of cochlear hair cell stereocilia tip links that bridge stereocilia and the widely accepted model for how mechanical forces generated by stereocilia movement are transduced into channel gating and hair cell depolarization^35–37^. In an attempt to gain a similar level of insight for tactile sensory neurons, we visualized the 3D architecture of the three Aβ RA-LTMR end organs in their native tissue context to identify ultrastructural features that may underlie mechanical gating and to ask whether these structural features are conserved across the different end organs. For this, we applied enhanced Focused Ion Beam Scanning Electron Microscopy (FIB-SEM)^38^ to image the three end organs in their entirety with 6-nm isotropic voxels (Extended Data Fig. 5A-C). Coupled with deep-learning based segmentation^39^, we fully reconstructed the Aβ RA-LTMR axons of the guard hair follicle lanceolate complex, the Meissner corpuscle, and the Pacinian corpuscle to generate fine 3D renderings of the three Aβ RA-LTMR end organs.

**Figure 5.**
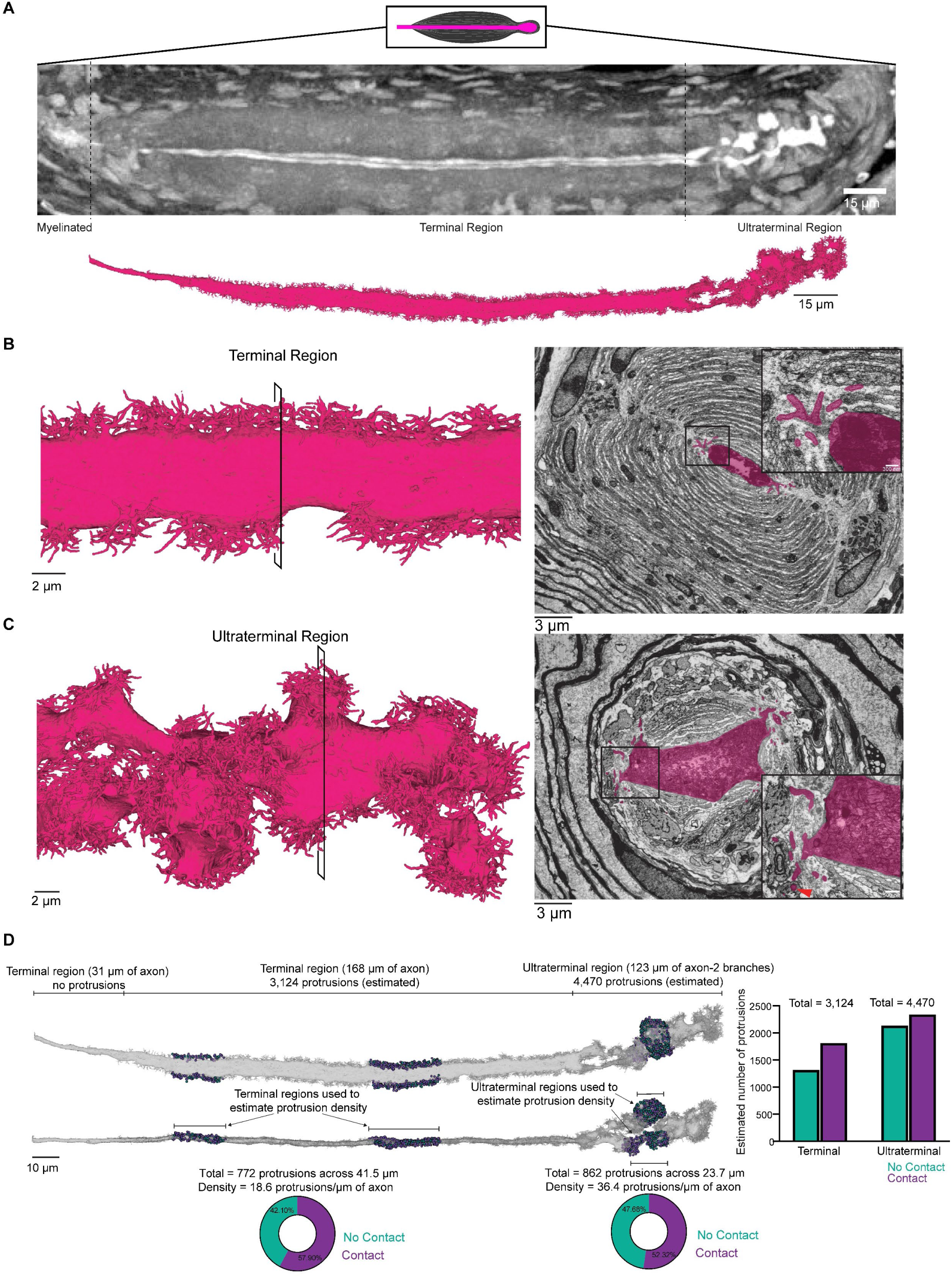
FIB-SEM reconstruction of a Pacinian corpuscle Aβ RA-LTMR reveals dense networks of axon protrusions that form contacts with corpuscle lamellar cells. (**A**) X-ray tomography of a Pacinian corpuscle (bright white axon) prior to FIB-SEM imaging (top) and corresponding 3D rendering of the Pacinian Aβ RA-LTMR after imaging (bottom). (**B**) Left, 3D rendering of a section of the Aβ RA-LTMR in the terminal region of the Pacinian corpuscle. Right, pseudo-colored image from the FIB-SEM volume with inset showing protrusions and their contacts with the non-neuronal lamellar cells of the inner core. (**C**) Left, 3D rendering of a section of the Aβ RA-LTMR in the ultraterminal region of the Pacinian corpuscle. Right, pseudo-colored image from the FIB-SEM volume with inset showing protrusions and their contacts with the non-neuronal lamellar cells of the inner and outer core. The red arrowhead shows an example of a protrusion encased by a lamellar cell at the boundary of the inner and outer core. (**D**) Characterization of protrusion terminals in the terminal and ultraterminal regions of the Pacinian corpuscle. Two representative regions in the terminal region and two representative regions in the ultraterminal region were selected to quantify and characterize protrusion terminals. The pie charts show the percentage of protrusions within the terminal or ultraterminal region that contact a lamellar cell (purple) or do not form a contact (teal). The statistics of protrusion terminals in these regions were used to estimate the total number of protrusions across the Pacinian corpuscle and whether those protrusions terminate by contacting a lamellar cell or within the collagen matrix within both the terminal and ultraterminal region (graphed on the right).

In hairy skin, we imaged a 5.12 x 10^5^ μm^3^ volume of tissue containing the entire Aβ RA-LTMR lanceolate complex surrounding a guard hair follicle, which is the longest hair type on the mouse trunk and accounts for 1% of all trunk hair follicles (Extended Data Fig. 5A). We identified and reconstructed all 47 lanceolate endings formed by heavily myelinated Aβ axons that encircle the outer root sheath of the hair follicle (Fig. 2A and Supplementary Video 1). We presume these to be the endings of two or three Aβ RA-LTMRs based on prior experiments assessing guard hair follicle innervation by genetically labeled LTMR populations^4, 40^. All but one axon branched at the base of the hair follicle complex to form multiple interdigitated lanceolate endings that extend within the longitudinal collagen matrix along the length of the hair shaft. In addition to the 47 Aβ RA-LTMR lanceolate endings, we were surprised to find six lanceolate endings formed by four unmyelinated, small caliber axons (Extended Data Fig. 6A). Smaller diameter Aδ- and C-LTMRs also form lanceolate endings associated with awl/auchene and zigzag hair follicles but not guard hair follicles^4^; however, CGRP antibody staining in hairy skin revealed CGRP^+^, NFH^-^ lanceolate endings (Extended Data Fig. 6B), suggesting that small diameter, CGRP^+^ fibers may form a small subset of lanceolate endings surrounding hair follicles. Each lanceolate ending was associated with processes originating from 17 TSCs, which were also embedded within the longitudinal collagen matrix and were fully reconstructed. These 17 TSCs were mostly non-overlapping, or tiled, and had cell bodies located at the base of the lanceolate complex (Fig. 2D, Supplementary Videos 2 and 3), consistent with previous reports^41^. Although the TSC processes formed an overlay that covered the lanceolate axons along their full extension adjacent to the hair shaft, intermittent gaps in TSC coverage formed at the TSC boundaries, leaving exposed small portions of axon membrane. These gaps most often formed on opposite sides of the lanceolate ending, along the axon membrane proximal and distal to the outer root sheath (Fig. 2A and Extended Data Fig. 6C). The membranes of these TSCs were studded with membrane pits that appear structurally analogous to the mechanoprotective, lipid-raft structure called caveolae found in several cell types, including skeletal and endothelial cells^42^. Indeed, deletion of the *Caveolin1* gene led to loss of these structures within TSCs of hairy skin (Extended Data Fig. 6C-D).

**Figure 6.**
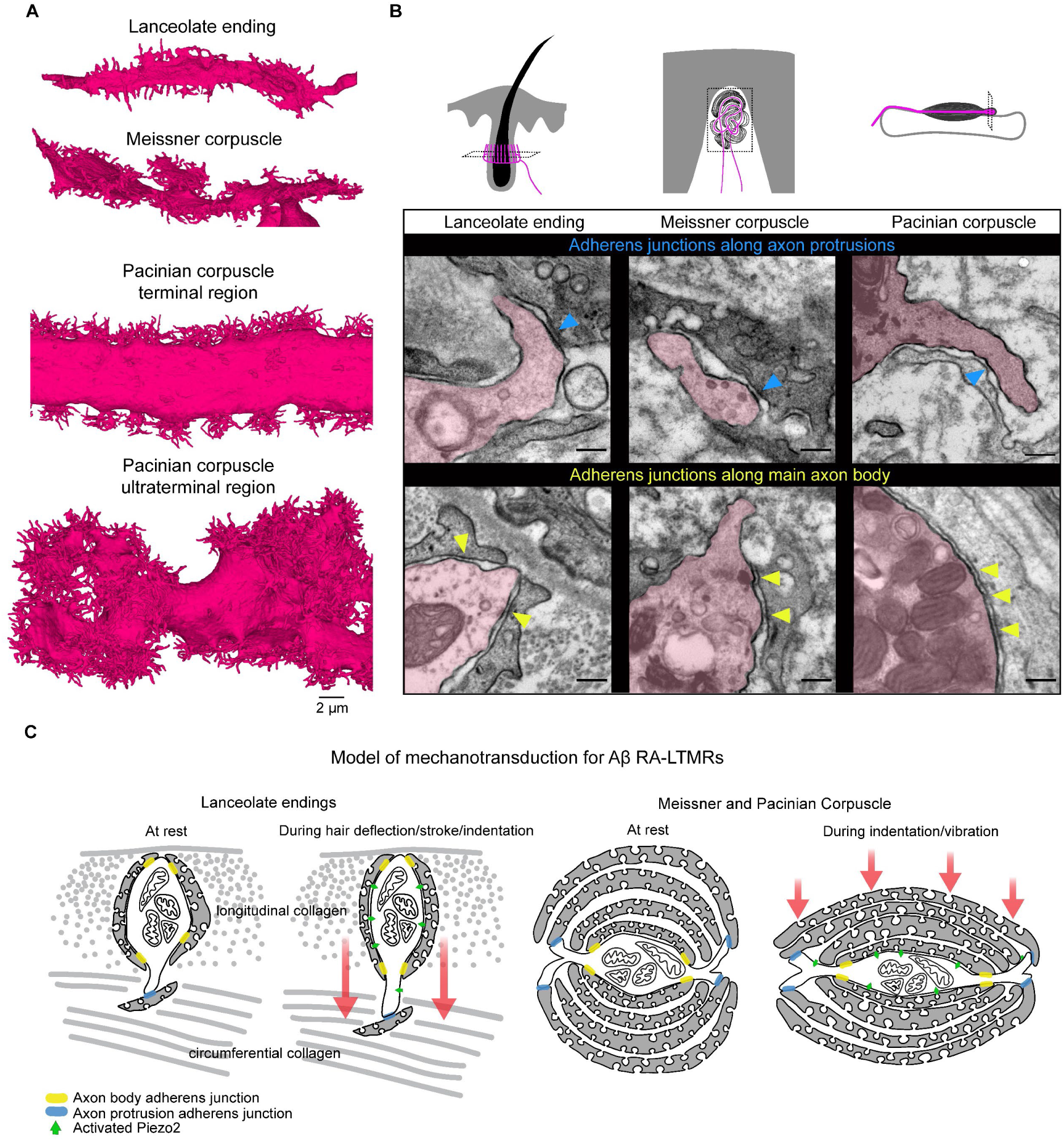
The conserved presence of adherens junctions at Aβ RA-LTMR axon protrusion contacts with non-neuronal cells across morphologically dissimilar mechanosensory end organs suggests a unified model of mechanotransduction. (**A**) 3D renderings of Aβ RA-LTMR axon terminals forming a single lanceolate ending, Meissner corpuscle afferent, and a portion of the terminal and ultraterminal region of the Pacinian corpuscle. Images are shown on the same scale. (**B**) Transmission electron microscopy (TEM) micrographs showing adherens junctions between the axon protrusions of Aβ RA-LTMRs and resident non-neuronal cells (blue arrowheads) and along the main body of the Aβ RA-LTMRs and neighboring terminal Schwann cells or lamellar cells (yellow arrowheads) in each end organ. This experiment was repeated in two animals. Scalebar, 200 nm. (**C**) Illustration of proposed model of mechanotransduction for Aβ RA-LTMRs. Adherens junctions at axon protrusions (blue) and along the main body of the Aβ RA-LTMR axon (yellow) serve as distinct anchor sites that render the Aβ RA-LTMR uniquely sensitive to dynamic stimuli. Hair deflection or skin indentation/vibration tugs on the hundreds to thousands of protrusions along the length of the axon within the end organ leading to stretching of the Aβ RA-LTMR membrane and activation of Piezo2.

Distal to the outer root sheath and longitudinal collagen network lies the circumferential collagen matrix that encircles the hair shaft at an orientation perpendicular to the longitudinal collagen network and contains the circumferential endings of the Aβ field-LTMRs and CGRP^+^ Aδ circumferential-high threshold mechanoreceptors (Aδ circ-HTMRs)^43–45^. These two circumferential sensory neurons are distinct from Aβ RA-LTMRs not just in their perpendicular orientation to one another, but also in their mechanical tuning. While Aβ field-LTMRs are sensitive to gentle stroking across the skin, both circumferential ending neuron types exhibit a higher activation threshold in response to skin indentation than Aβ RA-LTMRs^43, 44^. Within the volume, we identified and reconstructed two large caliber, myelinated axons we presume to be Aβ field-LTMRs, and we reconstructed a subset of circumferential TSCs that associated with these large caliber axons (Fig. 2B, Fig. 2E and Supplementary Video 4). We also reconstructed two small diameter axons within the circumferential collagen matrix that have the morphological properties of circumferential CGRP^+^ Aδ circ-HTMR endings (Fig. 2C, Extended Data Fig. 6E and Supplementary Video 5). The designation of the Aβ field-LTMRs and CGRP^+^ Aδ circ-HTMRs in our volume was further validated in parallel by genetically labeling each subtype with an EM reporter^46^ and identifying the structural homology by single section transmission electron microscopy (TEM) (Extended Data Fig. 6F). Indeed, by TEM, we observed abundant neurofilaments in Aβ field-LTMRs that were noticeably absent from small diameter CGRP^+^ Aδ circ-HTMRs, an ultrastructural distinction also observed between the two classes of circumferential endings in our FIB-SEM volume. In addition to the sensory axons and TSCs within the circumferential collagen, we found an extensive network of ∼17 non-neuronal cells, which we term circumferential support cells (CSCs), that reside in close proximity to the lanceolate endings. These previously uncharacterized CSCs are structurally distinct from TSCs in their minimal association with circumferential sensory axons, and their developmental origin and function in hair follicle physiology remain unclear. We reconstructed a subset of CSCs whose cell bodies reside in the circumferential collagen network distal to the lanceolate complexes and found that they fully encircle the guard hair in a vertically tiled manner (Fig. 2F and Supplementary Video 6).

To identify structural features underlying the Aβ RA-LTMR’s heightened sensitivity to mechanical forces, we examined features unique to the Aβ RA-LTMR lanceolate complexes. One strikingly unique ultrastructural feature of the Aβ RA-LTMRs was the presence of numerous axon protrusions that emerged only where the sensory axon formed close associations with TSCs (Fig. 3A-C), a location coincident with the axolemmal expression of Piezo2 (Fig. 1B and Extended Data Fig. 2A). These axon protrusions appeared along the length of the lanceolate endings at a density of 1.9 protrusions/µm of axon and extended through the intermittent gaps between TSC processes (Fig. 3B-C). Moreover, protrusions were absent from the TSC gaps proximal to the outer root sheath epithelial cells and were sparse or lacking in both the six small caliber lanceolate endings and the circumferential endings of Aβ field-LTMRs and CGRP^+^ Aδ circ-HTMRs (Extended Data Fig. 6A and 7A). Of the 1,742 axon protrusions originating from the 47 Aβ RA-LTMR lanceolate endings, 51% (880/1742) reached beyond the local longitudinal collagen matrix and extended into the circumferential collagen network (Fig. 3D and Extended Data Fig. 7B). Because of the potential for unique biomechanical properties under mechanical stress at the longitudinal/circumferential collagen matrix interface, we next examined the properties of the axon protrusions that extended across the interface and into the circumferential collagen region. We found that 57.3% (504/880) of these axon protrusions terminated by forming close associations with resident cells of the circumferential collagen matrix (Fig. 3D-E and Extended Data Fig. 7B). Furthermore, a significant proportion (32%, 163/504) of these contacts formed with the four reconstructed CSCs (Fig. 3E and Supplementary Video 7). In contrast, only a single contact was observed between the axon protrusions and the seven reconstructed circumferential TSCs (Extended Data Fig. 7C), revealing the preferential association between the axon protrusions and CSCs. These findings suggest that an extensive series of contacts between Aβ RA-LTMR lanceolate axon protrusions and CSCs serve to bridge the Aβ lanceolate endings across two perpendicularly oriented collagen matrices and anchor the sensory axon within the circumferential collagen region. These ultrastructural features suggest a model in which hair deflection or local skin indentation moves the circumferential collagen matrix relative to the longitudinal collagen matrix, thereby stretching Aβ RA-LTMR axon protrusions and lanceolate processes and gating Piezo2 localized along their membranes.

We next sought to determine whether similar ultrastructural features are prevalent in the other Aβ RA-LTMR end organs and, if so, whether a common model may explain mechanotransduction across them. Therefore, we imaged a 5.6 x 10^4^ μm^3^ volume of glabrous skin containing a Meissner corpuscle innervated by two myelinated Aβ LTMRs (Extended Data Fig. 5B and Supplementary Video 8). We reconstructed these two Meissner corpuscle Aβ LTMRs in addition to their associated lamellar cells as well as a subset of unmyelinated axons associated with the corpuscle, which may correspond to peptidergic and non-peptidergic C-fibers^47^ (Fig. 4A-C and Extended Data Fig. 8A). By confocal microscopy, the Meissner corpuscle appears as a spherical end organ with tortuous NFH^+^ axons (Fig. 1A). However, our reconstruction revealed two Aβ LTMR axons that traveled through the corpuscle in a largely linear manner (Fig. 4A-B), strikingly analogous in appearance to the hair follicle-associated Aβ RA-LTMR lanceolate endings (Fig. 3A). While the central Meissner Aβ axon was unbranched and shed its myelin at the base of the corpuscle structure, the more apical Aβ axon branched earlier within the dermal papilla and continued unmyelinated for 27.4 μm before entering the corpuscle (Fig. 4A). Four lamellar cells with cell bodies situated at the edge of the corpuscle extended numerous interdigitated lamellar processes to form the characteristic multi-layered wrappings of Meissner Aβ LTMRs (Fig. 4C). Three of the four lamellar cells contributed to the innermost wrapping of both Aβ LTMR axons, revealing a remarkable degree of interconnectedness of axons and lamellar cells within this structure (Fig. 4C). As with the TSCs of hairy skin, caveolae appear along the lamellar cell membrane; these were absent in mice lacking the *Caveolin1* gene (Extended Data Fig. 8B), revealing a conserved structural feature of TSCs of hairy skin and lamellar cells of Meissner corpuscles. Moreover, the lamellar cells had intermittent gaps in coverage along the sensory axon body (Fig. 4C and Extended Data Fig. 8C). These gaps often appeared at the vertices of the ellipsoid axon profile and their position remained consistent across the corpuscle. Strikingly similar to what we observed for hair follicle Aβ RA-LTMR lanceolate endings, an extensive array of axon protrusions emerged between these gaps and extended into the surrounding collagen network (Fig. 4C). These protrusions were restricted to the portion of axon within the corpuscle proper, suggestive of a role in mechanotransduction given their proximity to membrane-bound Piezo2 (Fig. 1C, Fig. 4B-C, Extended Data Fig. 2A, and Extended Data Fig. 3A).

Within most Meissner corpuscles exist two molecularly and physiologically distinct Aβ LTMRs^17^. The Meissner TrkB-expressing (TrkB^+^) population is the stereotypical, highly sensitive Aβ RA-LTMR, while the Meissner Ret-expressing (Ret^+^) subtype is, on average, less sensitive to skin indentation and exhibits varying rates of adaptation^17^. In addition to being molecularly and physiologically distinct, these two neuronal populations have distinct interactions with lamellar cells. Previous work detailing ultrastructural distinctions across the two Meissner afferent subtypes found that lamellar cell processes associated with the axons of TrkB^+^ Aβ RA-LTMRs were more elaborate and numerous than those associated with Ret^+^ Aβ LTMRs^17^. In our reconstruction, we found that across all depths of the corpuscle, the central axon (Fig. 4C, magenta) was wrapped more extensively by lamellar cell processes, suggestive of the central axon being TrkB^+^ and the peripheral axon being Ret^+^.

Using this classification for the two axons, we analyzed the axon protrusion density across these two distinct subtypes. We found that the putative TrkB^+^ Aβ RA-LTMR axon formed more than twice as many axon protrusions as the putative Ret^+^ axon (396:162, respectively) and had 11.7 protrusions/μm of axon—a density roughly 4 times that of the putative Ret^+^ axon (3 protrusions/μm), revealing a surprising second ultrastructural distinction for these two subtypes that may underlie their difference in tactile sensitivity (Fig. 4D). To test whether the density of axon protrusions along TrkB^+^ axons was consistent across corpuscles, we reconstructed the Aβ axon of a second, singly innervated Meissner corpuscle from a different FIB-SEM volume (Extended Data Fig. 8D). As at least 80% of corpuscles within the forepaw digits receive a TrkB^+^ afferent (Extended Data Fig. 8E) and as Meissner corpuscles are virtually absent in sensory neuron-specific TrkB knock-out mice^17^, we predict that the axon of this singly innervated Meissner corpuscle is TrkB^+^. We found that the presumed TrkB^+^ afferent of the singly innervated Meissner corpuscle was centrally located in the corpuscle and wrapped by numerous lamellar cell processes—features paralleled by the TrkB^+^ axon of the dual-innervated Meissner corpuscle (Extended Data Fig. 8D). Moreover, the axon of the singly innervated Meissner corpuscle had a density of 15.1 protrusions/μm of axon, a value comparable to the TrkB^+^ axon of the dual-innervated Meissner (Fig. 4D). In characterizing the terminal structure of the axon protrusions from both corpuscles, we observed a conserved structural motif analogous to Aβ RA-LTMR lanceolate endings. We found that 62% (334/538) and 67% (264/396) of protrusions from the presumed TrkB^+^ axon of the single and double-innervated Meissner corpuscles, respectively, contacted lamellar cell processes within the corpuscle (Fig. 4D and Supplementary Video 9). Although the density of protrusions was lower in the presumed Ret^+^ axon of the double-innervated Meissner corpuscle, a similar proportion of its protrusions formed intimate associations with lamellar cells (60%, 97/162) (Fig. 4D). Thus, the heightened density of axon protrusions in the more mechanically sensitive TrkB^+^ Aβ RA-LTMRs, compared to the Ret^+^ Aβ LTMR, suggests that these structures may contribute to the high sensitivity and low activation threshold of TrkB^+^ Aβ RA-LTMR axons of Meissner corpuscles.

The third end organ structure innervated by Aβ RA-LTMRs, the Pacinian corpuscle, is altogether unique in its size, location, simplicity in morphological appearance, and sensitivity to dynamic stimuli. Pacinian corpuscles of the mouse are mostly restricted to the joints and the periosteum surrounding bone, and are 200-300 µm in length and ∼40 µm in diameter^48, 49^ (Fig. 1A). Each Pacinian corpuscle is innervated by a single Aβ RA-LTMR axon that extends the length of the corpuscle, typically unbranched, before terminating in a branched, bulbous structure called an ultraterminal region^49^. Encircling the central axon are numerous concentric cellular processes^32^. The innermost rings that form the inner core around the axon are established by lamellar cells analogous to those of the Meissner corpuscle^50^. Although the Aβ LTMR of the Pacinian is analogous to the Aβ lanceolate ending and Meissner afferents in its RA response to static indentation, it is unique in its sensitivity to high frequency vibrations (>300 Hz)^51, 52^. To visualize the 3D ultrastructural features of the Pacinian Aβ RA-LTMR, we imaged a 9.6 x 10^5^ μm volume of tissue isolated from the periosteum membrane surrounding the fibula of a mouse and containing a Pacinian corpuscle (Extended Data Fig. 5C). We identified and reconstructed a single Aβ RA-LTMR axon that shed its myelin upon entering the corpuscle and remained unbranched within the terminal region for ∼200 µm before branching and entering the ultraterminal region (Fig. 5A and Supplementary Video 10). Within the terminal region, the cellular architecture remained simple and consistent. There, the Aβ RA-LTMR axon was ellipsoid (1.5 µm x 4.5 µm) with its major axis aligned with the longitudinal cleft (Fig. 5B), whose orientation was maintained through the terminal region. The multi-layered wrapping of the inner core region was established by ∼73 lamellar cells whose cell bodies were situated at the inner/outer core boundary (Extended Data Fig. 9A). The thin and densely packed lamellar cell processes emerging from the 73 lamellar cells precluded their reconstruction; however, we used *ROSA26-Confetti*, a stochastic, multicolor Cre-dependent fluorescent mouse reporter line, with a *Plp1-CreER* mouse to reveal the intermingled nature of lamellar cell processes that extend throughout Pacinian corpuscles—a blending of lamellar cell processes analogous to that observed within the Meissner corpuscle (Extended Data Fig. 9B). Intermittent gaps in lamellar cell coverage formed along the axon at the vertices of the major axis leaving the portion of axon membrane closest to the two longitudinal clefts exposed. As with the Aβ RA-LTMRs in hairy and glabrous skin, an extensive array of axon protrusions emerged from the central axon at these gaps and extended into the collagen matrix of the cleft (Fig. 5B). The structure of these axon protrusions was more complex than the Aβ RA-LTMRs of the hair follicle or Meissner corpuscle, resembling a tree with a trunk-like base from which numerous protrusions extended like branches. The base of these protrusions was often structurally supported by a dense, local collagen network that encircled the structure and disintegrated as the protrusions extended from the central axon and branched (Extended Data Fig. 9C). Because of the large size of the corpuscle and the density of protrusions within it, we characterized the protrusion density and terminal structure across four small regions of the volume and used these measurements to estimate the total number of protrusions across the corpuscle. We estimate there are ∼3,124 protrusions within the terminal region occurring at a density of 18.6 protrusions/µm of axon. We also estimate that ∼57.9% (1,809/3,124) of these protrusions terminated by contacting lamellar cell processes in the inner core (Fig. 5D), revealing a conserved structural motif analogous to the Aβ RA-LTMR lanceolate ending and Meissner corpuscle.

As the Pacinian’s Aβ RA-LTMR axon transitioned from the terminal to ultraterminal region, it branched and its structure became far more complicated. Within the ∼60 µm long ultraterminal region, which included 123 µm of axon, the axon expanded up to 12 µm in diameter and contained a remarkably high density of mitochondria (Fig. 5C). Accompanying the dissolution of the lamellar cell cleft was the emergence of axon protrusions of greater prevalence and structural complexity at many points along the circumference of the axon. The density of axon protrusions in the Pacinian’s ultraterminal region was estimated to be twice that of the terminal region, with 36.4 protrusions/µm of axon (estimated total = 4,470 protrusions). Across the two sampled regions of axon within the ultraterminal region, we found that 52.3% of protrusions terminated by contacting lamellar cell processes, suggesting that ∼2,339 protrusions within the ultraterminal region likely form intimate contacts with lamellar cells (Supplementary Video 11). Some of the terminals of the longer and more elaborate protrusions were encased by cells at the edge of the inner core or extended beyond the inner core to contact the innermost edge of the cells forming the outer core (Fig. 5C). The dense arrangement of axon protrusions within the ultraterminal region and their complex structure and reach across the inner and outer core boundary may impart elastic properties to the axon that facilitate its sensitivity to high frequency pressure transients and vibration across the entire corpuscle structure.

### Axon protrusions form cell junctions with non-neuronal cells across the three end organs

Collectively, from the full-scale end organ reconstructions we find that elaborate arrays of axon protrusions are a conserved ultrastructural feature across the highly sensitive Aβ LTMRs with rapidly adapting responses (Fig. 6A, reconstructions are displayed on the same scale). Indeed, the less sensitive neurons associated with the hair follicle and the Meissner corpuscle exhibit fewer protrusions (the Aβ field-LTMR, Aδ circ-HTMR, and Ret^+^ Meissner afferent) in comparison to their more sensitive counterparts (the Aβ RA-LTMR lanceolate ending and TrkB^+^ Meissner Aβ RA-LTMR). Furthermore, we found that across the three Aβ RA-LTMR end organs, the axon protrusions formed intimate associations with neighboring lamella or TSC processes, raising the possibility of physical contact points that tether the sensory axons to non-neuronal cells across the structure of each end organ. To further assess the ultrastructural nature of these intimate associations between axon protrusions and neighboring cells, we prepared high resolution TEM samples of each end organ using tannic acid, which enhances visualization of extracellular matrix components and identification of inter-cellular junctional complexes, such as adherins junctions^41^. Across the three end organs, we observed an abundance of fine filament-like structures that bridged the 15-20 nm gap between the sensory axons and their associated non-neuronal cells and spanned the membrane for 100-300 nm. These structures, which appeared structurally similar to adherens junctions, formed along the main body of the axon and neighboring TSC or lamellar cells, as has been observed previously in the hair follicles of rats^53, 54^, and on the axon protrusions at sites of contact with non-neuronal cells (Fig. 6B). Thus, axon protrusions not only greatly expand the axon surface area, and by extension the density of Piezo2 along the axolemma, but also anchor the sensory axon to non-neighboring cell processes via cell-cell junctions, which physically integrate the sensory axon into the end organ microenvironment.

## DISCUSSION

Our findings, which localize the requisite transduction channel Piezo2 to axonal membranes and identify tethered axon protrusions as a conserved structural motif shared across all Aβ RA-LTMR end organs, lead us to propose a unified model to explain Aβ RA-LTMRs’ responsiveness and entrainment to dynamic stimuli no matter the end organ structure they innervate. We propose that, like the hooks of Velcro or the burrs on the burdock plant, the protrusions of Aβ RA-LTMR axons (Fig. 6A) and their capacity to form cell-cell junctions act to fasten the sensory axon to distant cell processes thereby enhancing the bonding surface within the end organ and extending the reach of Piezo2 across the end organ structure. This conserved ultrastructural feature allows movement of hair or indentation or vibration of skin to stretch the axon membrane across hundreds to thousands of locations within an individual end organ structure, leading to Piezo2 gating and axonal excitation (Fig. 6C).

We found that Piezo2 localization in all three Aβ RA-LTMRs is restricted to terminal axons embedded within their respective end organs. The restricted localization of Piezo2 within the sensory axons and the lack of light-touch responses in neuronal Piezo2 knock-out animals^7, 8, 10, 11^, but not in *Dhh-Cre; Piezo2^f/f^* mice (Extended Data Fig. 4A-E), emphasizes the indispensable, cell-autonomous role of axonal Piezo2 in light touch; however, it remains possible that TSCs and lamellar cells express other mechanosensitive ion channels that modulate the tactile responses of Aβ RA-LTMRs. By reconstructing the end organ-innervating regions of axons containing Piezo2, we were able to characterize ultrastructural features unique to Aβ RA-LTMRs that may underlie their sensitivity to dynamic stimuli. Despite the dissimilar microenvironments in which each Aβ RA-LTMR resides—neighboring a hair follicle, associated with bone, or wedged within a dermal papilla near the surface of glabrous skin—we observed remarkable ultrastructural homology across these three end organ structures. We found an abundance of cytoplasmic axon protrusions to be a unique structural feature conserved across the Aβ RA-LTMRs within each end organ structure. Our observation that a large portion of axon protrusions formed by Aβ RA-LTMRs contact non-neuronal cells and form cell-cell junctions suggests a tether-like function for these axonal structures first described over 50 years ago^29–33^. Although our FIB-SEM volume did not contain slowly adapting (SA) Aβ LTMRs, which form the crown-like touch dome, previous ultrastructural analysis suggests that the terminal nerve plate of Aβ SA-LTMRs is smooth and lacks any discernable protrusion-like structures^19^. Collectively, this suggests these axon protrusions and their tethers are ultrastructural features reserved for the most sensitive sensory neurons involved in detecting dynamic touch.

Mechanical cell-cell coupling stabilizes tissue architecture and enables cells to sense and respond to tensile and shear forces in their microenvironment^55^. The abundance of junctions along the Piezo2-enriched region of Aβ sensory axons suggests that they play a central role in transforming mechanical stress across the end organ structure into sensory axon membrane strain, resulting in Piezo2 activation. We observed adherens junctions joining the main body of the sensory axon to its most proximate lamellar or TSC processes as well as at contact points between axon protrusions and more distant cell processes. We speculate that these junctions along the axon body and protrusions act as anchor points that enhance the transmission of forces from more peripheral portions of the end organ to the central axon body, helping to explain the low-threshold responses of Aβ RA-LTMRs. We found that in the hair follicle, ∼50% of protrusions from Aβ RA-LTMRs surrounding the guard hair extend beyond the longitudinal collagen network and enter the circumferential collagen matrix. Of the 880 protrusions that bridged these perpendicular collagen networks, over half form intimate contacts with cell processes embedded within the circumferential collagen matrix, perhaps tethering the protrusions within the circumferential collagen matrix and rendering the sensory axons uniquely sensitive to the shear, compressive, tensile, and torsional stress that occurs during hair deflection or nearby skin indentation. A majority of these contacts in the circumferential collagen network are made with a previously unidentified cell type, which we term the circumferential support cell or CSC, revealing a novel cellular component of lanceolate ending structure and possibly function. Previous work has shown that N-cadherin localizes to the cell-cell junctions that form between the sensory axons and TSCs of rat hair follicles^54^. Whether N-cadherin is required for the adherens junctions that form along the protrusions and within the circumferential collagen matrix described here, and the role of N-cadherin in mechanotransduction of Aβ RA-LTMRs remain to be explored.

In the Meissner corpuscle and Pacinian corpuscle, we speculate that tactile forces induce maximal bending and strain along the axon protrusions extending into the collagen network (Fig. 6C). These forces may be effectively transferred to the main body of the axon as a result of their cell-cell junctions formed between the extensive array of axon protrusions and the distant lamellar cell processes. In the Meissner corpuscle, we found that the more sensitive TrkB^+^ Aβ RA-LTMR exhibited twice the number of protrusions and four times the protrusion density of the less sensitive Ret^+^ sensory axon. While an equal proportion of protrusions from the two sensory neuron types formed contacts with lamellar cells within the corpuscle, we suspect that the higher density of protrusions of the TrkB^+^ axon render it more sensitive to tactile stimulation. Our reconstruction of the Pacinian corpuscle, and specifically of the ultraterminal region, revealed the densest array of axon protrusions among the Aβ RA-LTMRs. The tremendous enrichment of these structures may underlie the distinct tuning properties of this sensory axon. The Pacinian corpuscle is unique among LTMRs in its high-pass frequency tuning, potentially reflecting a unique resonance frequency of this end organ. In the mouse, these corpuscles are densest along mechanically conductive tissue, such as bone. We speculate that the dense mitochondria and the extensive axon protrusions and their lamellar cell contacts enable the Pacinian corpuscle to entrain to ultrahigh frequency mechanical stimuli conducted along the skeletal system of the extremities.

Collectively, our Aβ RA-LTMR end organ architecture and structural analyses reveal axon protrusions and their intimate contacts with resident, non-neuronal cells to be a common ultrastructural motif that we propose underlies the low force threshold activation of Aβ RA-LTMR axon membrane-bound Piezo2. We speculate that this motif exists in other sensory neurons equally sensitive to low-threshold, dynamic touch, including the Aδ- and C-LTMRs, which form lanceolate endings around zigzag and awl/auchene hairs^4^, and the sensory neurons of specialized end organ structures in the mucocutaneous skin of the lips, tongue, and genitalia.

Furthermore, given the structural homology of hair follicle lanceolate endings, Meissner corpuscles, and Pacinian corpuscles across species, we speculate that Aβ RA-LTMR end organ axon protrusions and the adherens junctions they form with surrounding support cells represents a basic functional unit of mechanotransduction in humans as well.

## METHODS

### Experimental model and subject details

Mice used in this study were maintained on mixed C57Bl/6J and CD1 backgrounds, except for mice used for the hair follicle (MH200121-B2K) and two Meissner Corpuscle FIB-SEM samples (200913FPT and 200913FPT2), which were pure C57BL/6J, and included both males and females. The *Piezo2^smFP-FLAG^*knock-in allele was generated at the Janelia Campus Research Gene Targeting and Transgenic Facility. The knock-in targeting construct was produced using recombineering techniques and traditional molecular cloning. A 5,197 bp of genomic DNA fragment containing exons 53 and 54 of the *Piezo2* gene was retrieved from BAC clone RP24-130D5. The *smFP-FLAG* gene was fused to Exon 54 followed by an frt-NeoR-frt cassette for ES cell selection. The homologous arms of the construct were 3,542 bp and 2,075 bp respectively.

To facilitate ES cell targeting Crispr/cas9 system was used. gRNA was invitro transcribed using MEGA shortscript T7 kit (Life Tech Corp AM1354). The template was PCR amplified using primers:

Forward5’- CCTTAATACGACTCACTATAGGCAAACTGATCACTTAAGACCTGGGTTTTAGAGCTA GAAATAGC

Revere 5’- AAAAGCACCGACTCGGTGCCACTTTTTCAAGTTGATAACGGACTAGCCTTATTTTAA CTTGCTATTTCTAGCTCTAAAAC

The gRNA was tested *in vitro* before being used for ES cell targeting. The test was carried out in a reaction of NEB 3.1 buffer 1ul, DNA template 120 ng, gRNA 40 ng, Cas9 160 ng in a total volume of 10 ul. The reaction mix was incubated at 37C for 2 hours. 0.5 ul proteinase K (20ug/ul) was then added and incubated at 57C for 30 min.

The targeting vector, cas9 protein (Fisher Scientific A36499 TRUECUT CAS9 PROTEIN V2) and the gRNA with concentrations of 10 ug, 3.75 ug and 1 ug respectively in total volume of 100 μl were co-electroporated into 1 million of G1 ES cells, which were derived from F1 hybrid blastocyst of 129S6 x C57BL/6J. Seventy-four G418 resistant ES colonies were isolated and screened by nested PCR using primers outside the construct paired with primers inside the construct. The primers used for ES cell screening were as following:

5’ arm forward primers: Sptbn1 Scr F1 (5’- CGCTCACTAGAGCAAAGTTG -3’) and Sptbn1 Scr F2 (5’- TAGGGTTCCTAGTAGGATCC -3’). Reverse primers: Cre scr R1 (5’– GAGGGACCTAATAACTTCGT -3’) and Cre scr R2 (5’- ATGATCGGAATTGGGCTGCA - 3’).

3’ arm forward primers: mMapple scr F1 (5’- CCATAGGATCGAGATCCTGT -3’) and mMapple scr F2, (5’- GACTACAACAAGGTCAAGCTGT -3’); Reverse primers: Sptbn1 Scr R1 (5’- CAGAGCAGCAGTTCTGACTT -3’) and Sptbn1 Scr 3R2 (5’-GACCCCAGAGATCTAATTCC -3’);

Thirty-eight of 48 ES cell clones screened contained both arms. Three of them were used for making chimeric mice. Chimeric mice were generated by aggregating the ES cells with 8-cell embryos of CD-1 strain. After achieving successful germline transmission of the *Piezo2^smFP-FLAG^* gene in one strain, the frt-NeoR-frt cassette was excised. Proper insertion of the gene into the Piezo2 locus was confirmed with loss of the wild-type Piezo2 gene in a mouse homozygous for the knock-in allele using the following primers:

Forward5’ Piezo2-Exon54: TGGAACTGGAGGAAGACCTCTACG

Reverse5’ Piezo2-smFP-FLAG: GAACAGCTCCTCGCCTTTCG

Reverse5’ Piezo2-3’UTR: CCCTGATTTCAGAGACATGGGAGT

To achieve specific labeling of sensory neuron subtypes, CreER driver lines were induced by administering tamoxifen dissolved in sunflower seed oil via intraperitoneal (I.P.) injection. Tamoxifen (MilliporeSigma) was dissolved in ethanol (MilliporeSigma) and then mixed with an equal volume of sunflower seed oil (MilliporeSigma). The mixture was vortexed, and ethanol was then removed under vacuum. The dosage for tamoxifen was as follows: for *Plp1^creER^*, mice received an I.P. injection of 1 mg of tamoxifen for 5 days (P15-P19); for *TrkB^CreER^*, mice, mice received an I.P. injection of 0.5 mg at P4; for *TrkC^CreER^* mice received an I.P. injection of 0.5 mg of tamoxifen at P5.

*Plp1^eGFP^* (JAX 033357)^56^ and *Plp1^CreER^* (JAX 005975)^57^ were used to label Schwann cells. *ROSA26^LSL-Matrix-dAPEX2^* (JAX 032765) and *ROSA26^FSF-Matrix-dAPEX2^* (JAX 032766) were used for genetic EM labeling^46^. *TrkC^CreER^* (JAX 030291) was used to label Aβ field LTMRs^43^. *CGRP^FleE^* was used to label Aδ HTMRs^58^. *Cav1^-/-^* (JAX 007083) was used to ablate caveolae^59^. *TrkB^CreER^* (JAX 027214)^60^ together with *Advillin^FlpO^*^58^ and ROSA26^FSF-LSL-tdTomato^ (JAX 021875)^61^ was used to selectively label Meissner Aβ RA-LTMRs. NPY2RGFP (GENSAT 011016-UCD) was used to label hairy Aβ RA-LTMRs. *Cdx2^Cre^*was used to delete Piezo2 in the sensory neurons and peripheral Schwann cells^62^. *Piezo2^fl/fl^* was used to delete Piezo2 in a Cre-dependent manner^20^. *Dhh^Cre^* was used to delete Piezo2 in peripheral Schwann cells (JAX 012929)^63^. Mice were handled and housed in standard cages in accordance with the Harvard Medical School and IACUC guidelines.

### Immunofluorescent staining

Tissue was isolated from P20 or older mice that were recently euthanized with isoflurane or sedated using a ketamine/xylazine mixture. Forepaw digit tips were cut to isolate tissue containing Meissner corpuscles. The back was treated with Nair to remove all hair and a large section of skin was cut. The fat on the underside of the hairy skin was removed to improve antibody penetration. Pacinian corpuscles were isolated from the periosteum membrane from the fibula of the mouse. The entire periosteum was isolated for whole mount staining. For staining of the Pacinian with cyrosections, *Plp1^eGFP^*mice were used to visualize and isolate Pacinian corpuscles in the periosteum membrane. Tissues were drop fixed in either 1% paraformaldehyde (PFA) in phosphate-buffered saline (PBS) for 1-2 hours at 4°C (for more sensitive antibodies) or Zamboni’s fixation buffer for 24 hours at 4°C. Following fixation, all tissues were rinsed in PBS and processed for 25 µm sections or whole mount staining as previously described^17^. For Extended Data Figure 8E, forepaw digit tips were sectioned at 200 µm and processed for whole mount staining. For each section, the proportion of Meissner corpuscles (labeled by S100B) that were innervated by a TrkB afferent (labeled by tdTomato) was calculated.

Primary antibodies were goat anti-mCherry (Sicgen, cat. no. AB0040-200, 1:500), chicken anti- GFP (Aves Lab, cat. no. GFP-1020, 1:500), rabbit anti-GFP (Invitrogen, cat. no. A-11122, 1:500), goat anti-GFP (US Biological, cat. no. G8965-01E, 1:500-1:1000), rabbit anti-S100 beta (ProteinTech, cat. no. 15146-1-AP, 1:300), rabbit anti-TUJ1 (Biolegend, cat. no. 802001, 1:500), chicken anti-neurofilament heavy chain (Aves, SKU: NFH, 1:300-1:500), rabbit anti-CGRP (Immunostar, cat. no. 24112, 1:375), and guinea pig anti-FLAG (1:500)^64^. The same laser settings and contrast/brightness adjustments were made for littermate controls of the *Piezo2^smFP-FLAG^* animals in Figure 1B and Extended Data Figure 2B.

### Electron microscopy sample preparation

Skin/tissue regions enriched for Meissner corpuscles, hair follicles, and Pacinian corpuscles were dissected as follows for electron microscopy (EM) sample preparations. The forepaw digit tips were dissected to isolate Meissner corpuscles. For hairy skin samples that focused on the ultrastructure of an isolated guard hair, all hairs except the guard hair were trimmed in a small piece of back hairy skin, allowing us to track the location of a guard hair follicle within the sample. For experiments that did not focus on guard hair ultrastructure, all hairs in the sample were trimmed. Pacinian corpuscles were isolated and dissected from the periosteum membrane surrounding the fibula using the fluorescent marker Plp1^eGFP^. Tissue samples containing forepaw digit tips, back hairy skin, and Pacinian corpuscles were immersed in a glutaraldehyde/formaldehyde fixative for 1 hour at room temperature, further dissected to remove muscle and fat, and subsequently fixed overnight at 4 °C. Sample preparation was done as previously described^46^. Ultrathin sections were cut at 50-70 nm and imaged using a JEOL 1200EX transmission electron microscope at 80 kV accelerating voltage. Images were cropped and adjusted to enhance contrast using Fiji/ImageJ. For tannic acid treatment, samples were incubated in cacodylate buffer containing 1% low molecular weight tannic acid for 30 min (Electron Microscopy Sciences) between the osmication step and the uranyl acetate step, with washes preceding and following this treatment. Ultra-thin sections were then stained with uranyl acetate and lead citrate. All single section TEM images presented in the paper were pseudo-colored by hand.

### FIB-SEM sample preparation

Skin/tissue regions enriched for Meissner corpuscles, hair follicles, and Pacinian corpuscles were dissected as described above. Four durcupan embedded mouse end organ samples: one hair follicle sample (MH200121-B2K), two Meissner Corpuscle samples (200913FPT and 200913FPT2), and one Pacinian Corpuscle sample (Pacinian2A) were prepared for FIB-SEM as described previously^65^. Specifically, each sample was first mounted to the top of a 1 mm copper post, which was in contact with the metal-stained sample for belter charge dissipation, as previously described^38^. The vertical sample posts were each trimmed to a small block containing the Region of Interest (ROI) with a width perpendicular to the ion beam, and a depth in the direction of the ion beam. The block sizes were 105 x 100 µm^2^, 90 x 70 µm^2^, and 85 x 75 µm^2^, and 140 x 120 µm^2^ for MH200121-B2K, 200913FPT, 200913FPT2, and Pacinian2A respectively. The trimming was guided by X-ray tomography data obtained by a Zeiss Versa XRM-510 and optical inspection under a Leica UC7 ultramicrotome. Thin layers of conductive material of 10-nm gold followed by 100-nm carbon were coated on the trimmed samples using a Gatan 682 High-Resolution Ion Beam Coater. The coating parameters were 6 keV, 200 nA on both argon gas plasma sources, and 10 rpm sample rotation with 45-degree tilt.

### FIB-SEM 3D large volume imaging

These four FIB-SEM prepared samples, MH200121-B2K (guard hair follicle volume), 200913FPT (single innervated Meissner corpuscle), 200913FPT2 (dual innervated Meissner corpuscle) and Pacinian2A (Pacinian corpuscle) were imaged by four customized Zeiss FIB-SEM systems previously described^38, 66, 67^. Each block face of ROI was imaged by a 1-nA electron beam with 0.9 keV landing energy. The x-y pixel resolution was set at 6 nm. A subsequently applied focused Ga+ beam of 15 nA at 30 keV strafed across the top surface and ablated away 6 nm of the surface. The newly exposed surface was then imaged again. The hair follicle (MH200121-B2K) and Meissner Corpuscle (200913FPT and 200913FPT2) samples were imaged at 1 MHz throughout their entire volumes. The ablation – imaging cycle continued about once every three and a half minutes for five weeks to complete FIB-SEM imaging MH200121-B2K, and about once every minute for one week to complete 200913FPT and 200913FPT2. To best balance the total acquisition duration and the image quality (signal to noise ratio) in critical sections, the Pacinian Corpuscle sample was imaged sequentially using three different SEM scanning rates of 3 MHz, 2 MHz and 1 MHz for the top (myelinated region and part of the terminal region), middle (terminal region and part of the ultraterminal region) and bottom (ultraterminal region) sections of ROI, respectively. The ablation – imaging cycle continued about once every 50 seconds for 16 days to complete the top section ROI of 55 x 60 x 185 µm^3^, continued about once every 65 seconds for 9 days to complete the middle section ROI of 55 x 60 x 70 µm^3^, and continued about once every 2 minutes for 13 days to complete the bottom section ROI of 55 x 60 x 60 µm^3^. The entire Pacinian2A sample was FIB-SEM imaged for 38 days.

Each acquired image stack formed a raw imaged volume, followed by post processing of image registration and pairwise alignment using a Scale Invariant Feature Transform (SIFT) based algorithm. The aligned stack consists of a final isotropic volume of 80 x 80 x 80 µm^3^, 30 x 40 x 50 µm^3^, 35 x 40 x 40 µm^3^ and 55 x 60 x 291 µm^3^ for MH200121-B2K, 200913FPT, 200913FPT2, and Pacinian2A respectively. The voxel size of 6 x 6 x 6 nm^3^ was maintained for each sample throughout the entire volume, which can be viewed in any arbitrary orientations. 174 nm was missing in MH200121-B2K between slice 6483 and slice 6484 due to microscope issues.

### Global image alignment and processing

Three slices (7597, 7598, 7962) were discarded in MH200121-B2K due to microscope issues, and adjacent slices (7596, 7599, 7961) were copied over. For global alignment, the SIFT-aligned FIB-SEM stacks were downsampled 32x to 192 nm x 192 nm x 192 nm voxel size and adjusted for contrast to reduce illumination unevenness. μCT volumes were cropped to just large enough to fully include the FIB-SEM ROI. The downsampled FIB-SEM stacks were used as moving images to align to μCT stacks which were fixed images using elastix^68, 69^. The elastix alignment was done in a manner similar to Phelps et al. (2021)^70^ (https://github.com/htem/run_elastix), where an affine alignment was followed by a B-spline elastic alignment. Mutual information (AdvancedMattesMutualInformation) was used as the main metric, and 28 grid spacing and 0 bending weight was used for B-spline alignment to avoid distortions. Corresponding points were added whenever necessary. After satisfactory alignment was achieved, the transform was inverted to allow the mapping of coordinates in the SIFT-aligned FIB-SEM space to the μCT space. In order to align images by translation alone to reduce distortion, the geometric center of each section was mapped to the μCT space, given that it is rotationally invariant and in general in a well aligned region for elastix alignment. Since the z-axes were nearly identical in direction for the SIFT-aligned FIB-SEM volume and the μCT volume, these transformed coordinates can then be used as displacement vectors to align raw image stacks.

To generate the final volumes, raw FIB-SEM images were first processed to clean up milling artifacts using Fourier transform^38^, and then contrast enhanced using CLAHE. Images were then placed using displacement vectors generated above into an aligned volume. Volumes were rotated in 3-D space to align their z-axes to major anatomical axes, such as the hair shaft, for the ease of analysis

### Automated segmentation and reconstruction

We used Segway, a segmentation pipeline previously developed for segmentation of transmission EM data^39^ and later refined for X-ray and isotropic data segmentation^71^, to automatically segment the FIB-SEM datasets. Segway uses a 3D U-Net convolutional neural network (CNN) model to create an affinity map to predict a ‘connectedness’ probability of each voxel to adjacent voxels^72^. CNN models were trained using ground truth labels generated through manual segmentation of small image volumes that included the sensory axons, non-neuronal cells of interest, as well as "background" regions (e.g., "empty" areas that should be masked out for reconstruction) using webKnossos^73^. A separate CNN model was generated for each dataset using volume-specific ground truth for MH200121-B2K (guard hair follicle volume), 200913FPT2 (dual-innervated Meissner corpuscle volume), and Pacinian2A (Pacinian corpuscle volume), except for 200913FPT which used the same CNN model as 200913FPT2 due to their homology. For CNN training we used Gunpowder^74^ to randomly sample batches and perform data augmentation; the network architecture and training parameters are as previously described^71^, though in these models we also use long-range affinities^75^ to improve prediction accuracy. After the models were trained, Daisy^76^ was used to deploy the CNN models and subsequent post-processing steps in the Segway pipeline to generate the output segmentation.

All volumes were segmented at 6 x 6 x 6 nm resolution except the Pacinian volume which was segmented at 12 x 12 x 12 nm to increase performance speed without a noticeable increase in error. To proofread and reconstruct neurons, we used MD-Seg, a merge-deferred segmentation proofreading method^39^. This method pre-agglomerates segmentation fragments only in a local block instead of across the entire volume to avoid catastrophic merge error propagations that often occur in large volumes. Inter-block merge decisions are computed and accessible to the proofreader during the reconstruction in the user interface using a hotkey. This pre-agglomeration threshold was 0.5 (out of a range from 0.0 to 1.0) for all volumes except for Pacinian2A which was 0.9 because only the axon was constructed and to minimize proofreading time of split errors.

To reconstruct the sensory neurons and non-neuronal cells of interest within our FIB-SEM volumes, we first identified the cells to be reconstructed based on their stereotypical morphological characteristics and selected a cell fragment that was contained within a single block. Using the hotkey, mergeable fragments in adjacent blocks were added to the single block from all sides. Sequential use of the hotkey enabled the proofreader to ‘grow’ the cell by continually adding computed merge fragments at the boundaries of the growing cell. In the event of a merge error, growth from the block containing the error could be blocked and removed from the reconstruction to prevent any further growth from the merged segment.

In each volume, all myelinated Aβ sensory neuron axons were reconstructed. Also, a subset of small diameter neurons that innervated the hair follicle and Meissner corpuscle were reconstructed. The terminal Schwann cells that associated with the lanceolate endings in the MH200121-B2K (guard hair follicle volume) and the lamellar cells that associated with the Aβ sensory axons in the 200913FPT2 (dual-innervated Meissner corpuscle volume) were reconstructed to visualize the intimate interactions between the non-neuronal cells of the end organs and the Aβ sensory axons. Additionally, we reconstructed a subset of the circumferential terminal Schwann cells that associated with the Aβ field LTMRs and a subset of previously uncharacterized circumferential support cells (CSCs) because of their numerous contacts with the protrusions of the Aβ sensory axons of the hair follicle. Based on structural homology, we estimate there to be roughly 17 total CSCs with cell bodies that reside in the circumferential collagen matrix on the same horizontal plane as the lanceolate endings. Additional non-neuronal cells were observed in the volume but were not reconstructed. All single sections from FIB-SEM volumes presented in the paper are pseudo-colored by the trained networks.

### Immuno-electron microscopy sample preparation

Skin regions enriched for Meissner corpuscles, hair follicles, and Pacinian corpuscles were isolated from *Piezo2^smFP-FLAG^* mice or littermate controls. The *Plp1^eGFP^* allele was also present in all mice to aid sample preparation. Forepaw digit tips were isolated as described above. For back hairy skin, all hairs were trimmed short, and a rectangular piece of skin (∼1 mm x 2 mm) was dissected with the rostral/caudal axis oriented along the long axis of the sample. Pacinian corpuscles were isolated as described above. Samples were drop fixed in 4% PFA in 0.1 M phosphate buffer (PB) for 2 hours at room temperature. Samples were micro-dissected 1 hour into fixation to remove access fat and muscle and returned to fix for the remainder of the 2 hours.

After samples were washed in 0.1 M PB, they were cryoprotected in 30% sucrose in 0.1 M PB. Samples were then embedded in Neg-50 Frozen Section Medium (VWR, cat. no. 21008-918). Immediately after the samples were embedded, they were cryosectioned into 50 µm sections and placed into chilled 0.1 M PB. Plp1^eGFP^ mice were used to visualize and isolate sections with end organ structures. Sections were then incubated in 50 mM glycine in 0.1 M PB for 30 minutes.

After aldehyde quenching, sections were blocked in 10% normal goat serum (Vector labs, cat. no. S-1000), 0.5% fish gelatin (Sigma, cat. no. G7041), and 0.05% Triton X-100 in 0.1 M PB for 2 hours at room temperature. Sections were then incubated with primary antibody in a modified blocking solution—10% normal goat serum, 0.5% fish gelatin, 1:100 guinea pig anti-FLAG^64^ in 0.1 M PB—for 24 hours at room temperature under gentle agitation. After a series of washes in 0.1 M PB, sections were incubated overnight with species-specific gold-labeled secondary antibody in a modified blocking solution—10% normal goat serum, 0.5% fish gelatin, 1:50 Nanogold-Fab’ goat anti-guinea pig IgG (Nanoprobes, cat. no. 2055) in 0.1 M PB. After washes in 0.1 M PB, sections were post-fixed with 1% glutaraldehyde for 10 minutes before being thoroughly rinsed with dH2O.

To reveal gold labeling, sections were incubated with HQ Silver Enhancement (Nanoprobes, cat. no. 2012) for 8 minutes under gentle agitation. The enhancement reaction was quenched by the addition and subsequent rinses with dH2O. Following silver staining, samples were stained with 1% osmium tetroxide in 0.1M PB for 30 minutes and subsequently 1% uranyl acetate in 0.05 M maleate buffer overnight at 4°C. Sections were then dehydrated with serial ethanol and propylene oxide (VWR, cat. no. 20411) dilutions before infiltration and embedding in an epoxy resin (LX-112, Ladd Research) mix and cured at 60°C for 48-72 hours. Ultrathin sections were cut at 50-70 nm and imaged using a JEOL 1200EX transmission electron microscope at 80 kV accelerating voltage. Images were cropped and adjusted to enhance contrast using Fiji/ImageJ.

### Protrusion ending quantification

To characterize the terminal properties of axon protrusions, the final 50% of a protrusion was examined to assess whether it formed a contact (<30 nm of space between membranes) with a neighboring cell. In the hair follicle volume, protrusions along the Aβ RA-LTMRs were categorized as terminating either in the local longitudinal collagen matrix or in the more distal circumferential collagen network. If a protrusion extended into the circumferential collagen network and came within ∼30 nm of a terminal Schwann cell that was also within the circumferential collagen, then it was counted as a contact with a cell in the circumferential collagen. Protrusions were also counted along the Aβ field-LTMRs. Small bumps along the sensory axons were not counted as protrusions. The same metrics were used to characterize the axon protrusions within both Meissner corpuscle FIB-SEM volumes. Because of the large size of the Pacinian corpuscle and the density of protrusions, we counted the number of protrusions and characterized their terminal endings in representative regions of the corpuscle and used that quantification to estimate the total number of protrusions and their terminal structures within the terminal and ultraterminal region. Two different regions in the terminal and ultraterminal region were selected to characterize protrusions to ensure we captured as much heterogeneity in protrusion density and terminal structure as possible across this large structure.

### *In vivo* DRG electrophysiology

Juxtacellular recordings were made from DRG neurons *in vivo*. Mice of both sexes were anesthetized with urethane (1.5 g/kg). An incision was made over the lumbar spine and scalp. A headplate was fixed to the skull using cyanoacrylate adhesive. The muscle overlying the DCN was removed, and the dorsal aspect of the C1 vertebrae was removed. The L4 DRG was exposed and two custom made spinal clamps fixed the spine in place. A platinum-iridium bipolar stimulus electrode (FHC) was lowered into the caudal DCN or C1 DC. A glass electrode (borosilicate, 1-2 MO) filled with saline was lowered into the DRG. Electrical stimuli (25-300 µA, 0.2 ms duration) were delivered to the DCN/DC at 0.5-1 Hz while searching for a unit. Once an antidromically activated unit was isolated, the receptive field was determined by brushing the hindlimb and thigh. If no receptive field could be identified, a series of stimuli were systematically delivered to the hindpaw and thigh to search for activation. Once the receptive field was located, one of three stimuli were delivered. Airpuffs (5 PSI from a 20 gauge needle, 1-3 mm from the hair) were used for hairy skin. For glabrous skin, a custom-built mechanical stimulator delivered static indentations (0-25 mN) using a 0.2 mm diameter probe tip. In some cases, vibration (10-1000 Hz, 0-25 mN) was delivered using the same stimulator. Signals were amplified using a Multiclamp 700A/B and acquired using a Digidata 1550A/B using Clampex 10/11 software. For Aβ RA-LTMRs shown in Extended Data Fig. 1A, cells were visualized under a microscope in BAC-transgenic *NPY2R-GFP* mice^4^ and large glass electrodes (20-30 µm tip diameter) were used to perform targeted recordings as previously described^17^. Force indentations (0.5 s duration 1-75 mN intensity) and sine vibrations of varying frequency and intensity were used to characterize the response properties of five NPY2RGFP^+^ cells isolated from four animals. Similar responses were observed for all isolated cells.

### DRG electrophysiology analysis

Units that were antidromically identified from DCN/DC stimulation were sorted into categories based on their response properties. Units that had their mechanical receptive field on hairy skin and were activated by movement of hairs were classified as ‘hairy’. Units that had receptive fields on glabrous skin were classified as ‘glabrous RA/SA’. Units that did not have a cutaneous receptive field but could become phase-locked to high frequency vibration (>300 Hz) were classified as Pacinian. Units were also detected that specifically responded to mechanical stimulation of the toenail. For Extended Data Fig. 4 these units were combined with pacinians as they are associated with bone and phase-locked to high frequency vibration (300 Hz). Units that lacked a specific cutaneous receptive field, but responded strongly to bending of the toe were identified as ‘proprioceptors’. If units in any animal did not respond to airpuff, brush, or strong vibration delivered to the thigh, trunk, or hindlimb, the units were classified as ‘insensitive’.

In *Cdx2^Cre^; Piezo2^Fl/Fl^* animals receptive fields were not discernable by brushing. Airpuffs and brushes were delivered systematically across the hindlimb and thigh. The hindlimb was pinched if no mechanically evoked responses could be found in order to confirm that the nerve from the DRG to the periphery was not damaged. Pinching in some cases evoked firing in antidromically-identified units, but in many recordings activated units that were not activated by DCN/DC stimulation.

### Data availability

All data, including FIB-SEM volumes and 3D models, generated during this study will be available by request.

### Code availability

Custom scripts used in this study will be available by request.

## Supporting information

Supplementary Table of Contents

## ACKNOWLEDGEMENTS

We thank Maria Ericsson, Giovanni De Nola, and Shachar Dagan for advice and help with electron microscopy preparations. We thank Caiying Guo from the Gene Targeting and Transgenics Facility at Janelia Research Campus for generating the *Piezo2^smFP-FLAG^* mouse. We thank Harald F. Hess for early input on this project. We thank Soha Ashrafi, Alan Emanuel, Vanessa Ruta and members of the Ginty lab for comments on the manuscript. This work was supported by NIH grants NS97344 and AT011447 (DDG), MH117808 (WCAL), the Edward R. and Anne G. Lefler Center for Neurodegenerative Disorders (DDG), a Howard Hughes Medical Institute–Jane Coffin Childs Fellowship (AH), the Howard Hughes Medical Institute (SP and CSX), and Stuart H.Q. & Victoria Quan Fellowship (QZ). Portions of the work were supported by a Foundry Award for the HMS Connectomics Core. DDG is an investigator of the Howard Hughes Medical Institute. This article is subject to HHMI’s Open Access to Publications policy. HHMI lab heads have previously granted a nonexclusive CC BY 4.0 license to the public and a sublicensable license to HHMI in their research articles. Pursuant to those licenses, the author-accepted manuscript of this article can be made freely available under a CC BY 4.0 license immediately upon publication.

## AUTHOR CONTRIBUTIONS

AH, QZ, SP, CSX, and DDG conceived the study. AH and QZ, with the help of MI, MNT, and SC, prepared all EM samples. Following resin embedding of FIB-SEM samples, SP prepared all FIB-SEM samples for imaging. FIB-SEM sample imaging was performed by SP and CSX. Post data registration and supervision of FIB-SEM samples was performed by CSX. Image data processing was performed by QZ and AH. Network training and automatic segmentation was performed by TMN under the supervision of WCAL. Ground truth for FIB-SEM volumes was annotated by RP, BS, KA, AS, BB, MK, NA, SC, AH, and QZ. AH, MI, and MNT performed immunohistochemistry and immuno-EM experiments. Proofreading of the models was performed by AH, QZ, and MNT. MNT quantified axon protrusions. AH performed the physiology experiments in Extended Data Figure 1A. JT performed electrophysiology experiments in Extended Data Figure 1B-C and Extended Data Figure 4. WX, ECP, and SC performed preliminary experiments related to reconstructions of FIB-SEM volumes, protrusion quantifications, and immuno-EM experiments. GR performed preliminary electrophysiology experiments. CS performed preliminary histology experiments. AH and DDG wrote the paper with input from all authors.

## EXTENDED DATA FIGURES

**Extended Data Figure 1.**
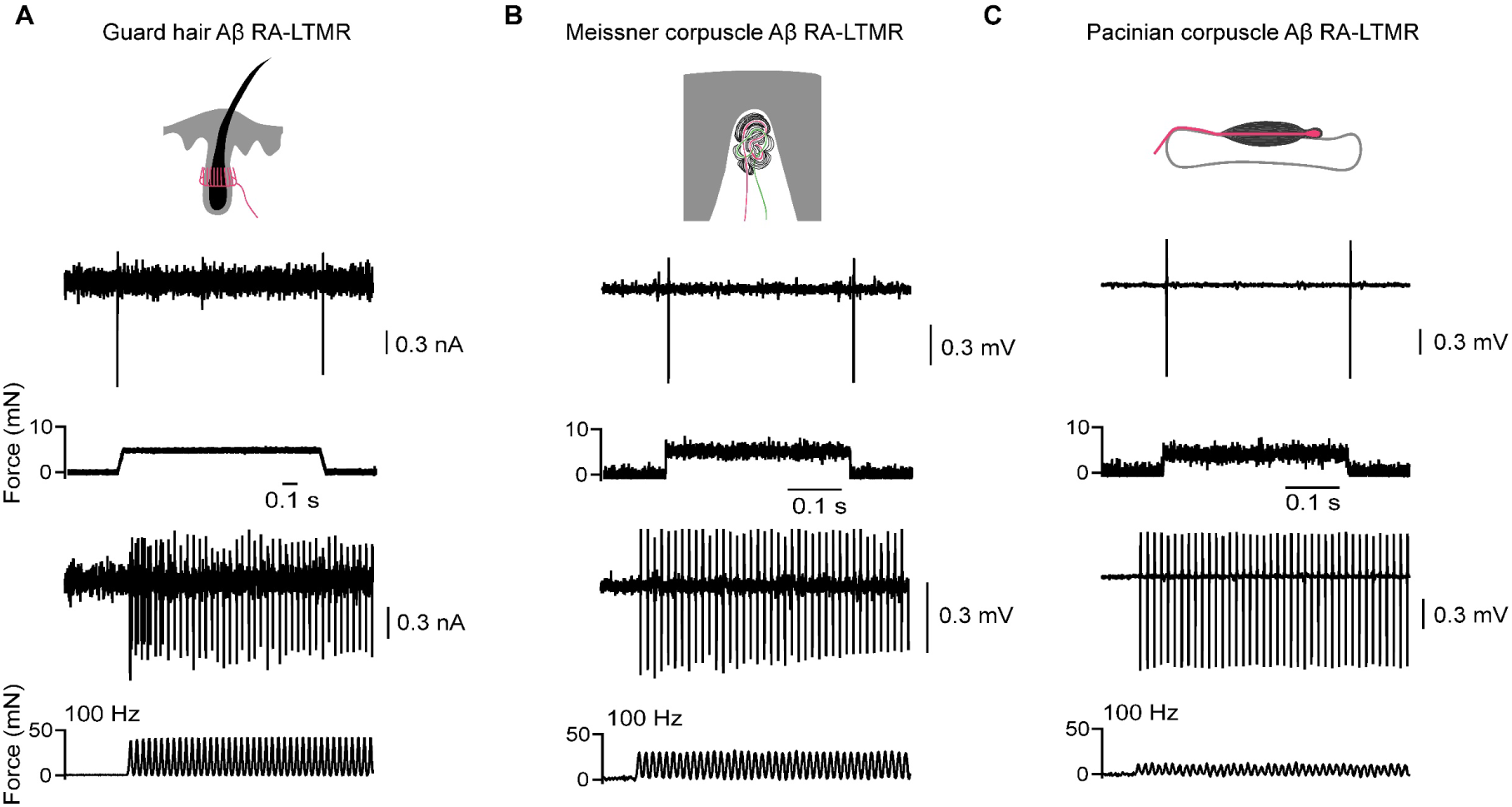
Aβ RA-LTMRs innervating the hair follicle, Meissner corpuscle, and Pacinian corpuscle all encode dynamic stimuli. (**A**) In vivo recording of an Aβ RA-LTMR forming lanceolate endings from an *NPY2R-GFP* mouse in response to step indentations and 100 Hz vibration delivered to the center of the receptive field of hairy thigh skin. Similar responses were observed in five GFP+ cells of four *NPY2R-GFP* mice. (**B**) In vivo recording of a representative Aβ RA-LTMR unit innervating a Meissner corpuscle in response to step indentation and 100 Hz vibration delivered to the center of the receptive field on glabrous hindpaw. (**C**) In vivo recording of a representative Aβ RA-LTMR unit innervating a Pacinian corpuscle in response to step indentation and 100 Hz vibration delivered to the ankle.

**Extended Data Figure 2.**
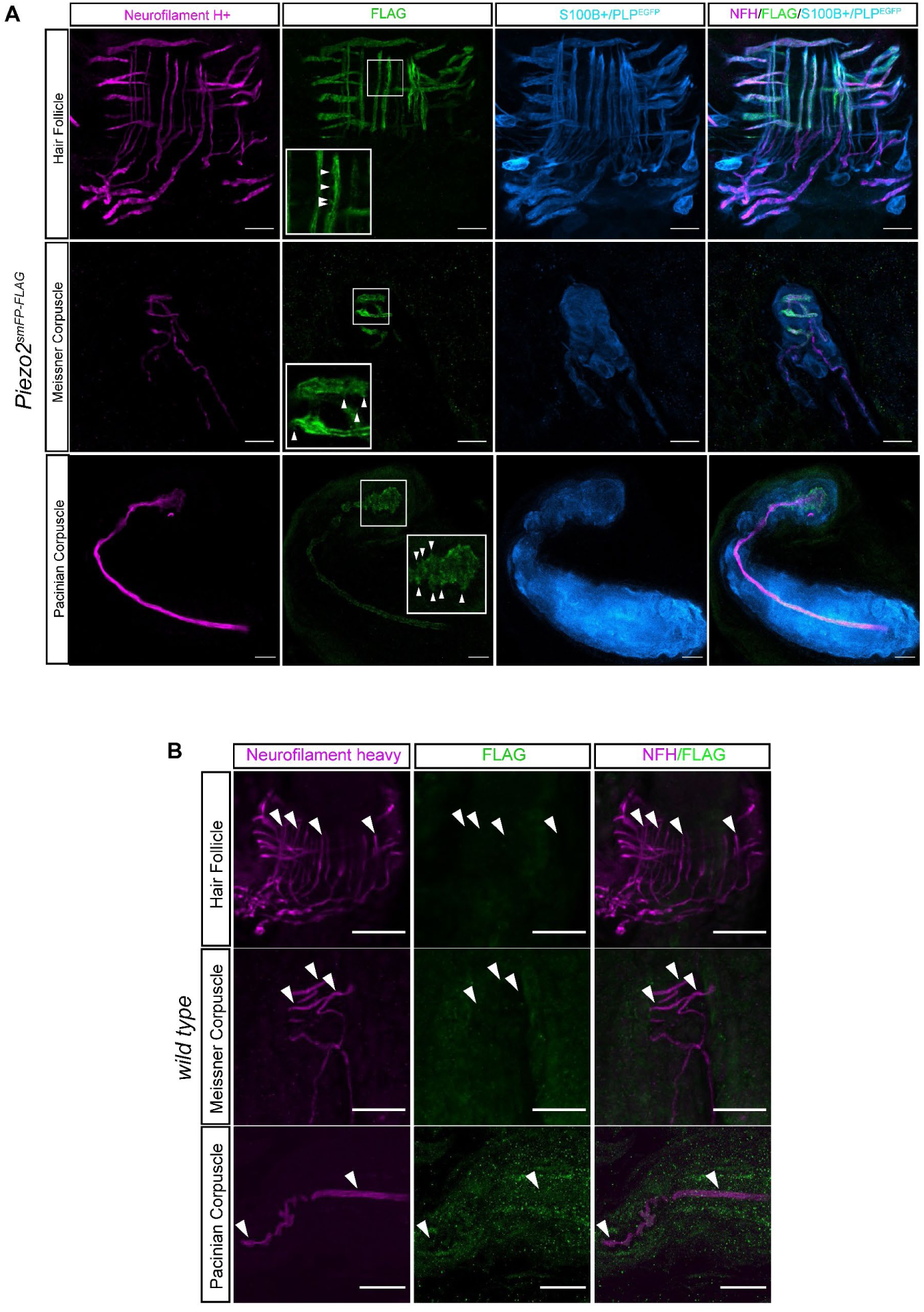
Piezo2-FLAG localizes to protrusion-like processes of Aβ RA-LTMRs and is not observed in control animals. (**A**) Piezo2-FLAG protein localizes to protrusion-like processes along the NFH^+^ Aβ RA-LTMRs forming lanceolate endings in hairy skin, Meissner corpuscles in glabrous skin, and the Pacinian corpuscle (white arrowheads). Piezo2-FLAG is not present within the cell bodies of TSC or lamellar cells (located at the base of the hair follicle and Meissner corpuscle, respectively) and is not observed above background tissue levels in the spherical cloud of lamellar cell processes within the Meissner corpuscle (S100B^+^) and Pacinian corpuscle (PLP^EGFP^). Scalebar, 10 µm. (**B**) FLAG and NFH co-staining in wild-type littermates of Piezo2^smFP-FLAG^ animals performed side-by-side and processed with same confocal settings as Figure 1B. White arrowheads show location of NFH^+^ Aβ RA-LTMRs forming lanceolate endings and innervating the Meissner corpuscle and Pacinian corpuscle. Scalebar, 25 µm.

**Extended Data Figure 3.**
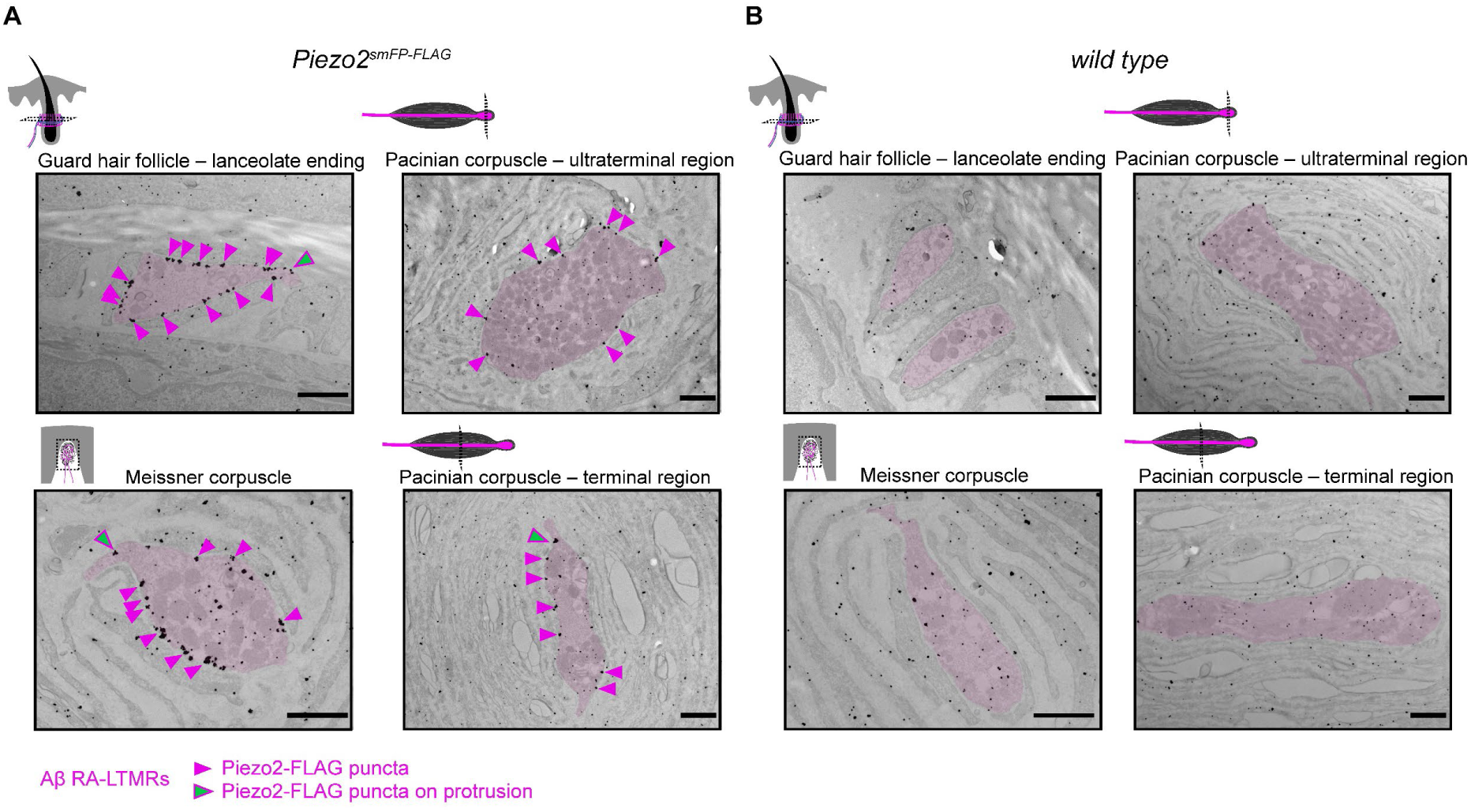
Additional examples of Piezo2 enrichment along the membrane of Aβ RA-LTMRs. (**A**) Immuno-electron micrographs from a second *Piezo2^smFP-FLAG^* animal stained for FLAG and processed with silver enhancement show enriched expression of the Piezo2-FLAG fusion protein along the axon membranes of the Aβ RA-LTMRs that form lanceolate endings around the hair follicle and innervate the Meissner corpuscle and Pacinian corpuscle within both the terminal and ultraterminal region (Aβ RA-LTMRs pseudo-colored pink). Piezo2-FLAG puncta can be seen along the sensory axon membranes (pink arrowheads) and along small cytoplasmic axon protrusions (green arrowheads). Not all puncta are labeled by arrowheads. Scalebar, 1 µm. (**B**) Immuno-electron micrographs from a wild-type littermate of the *Piezo2^smFP-FLAG^* animal shown in (A) stained for FLAG and processed with silver enhancement. The Aβ RA-LTMRs of each end organ are pseudo-colored pink. Scalebar, 1 µm.

**Extended Data Figure 4.**
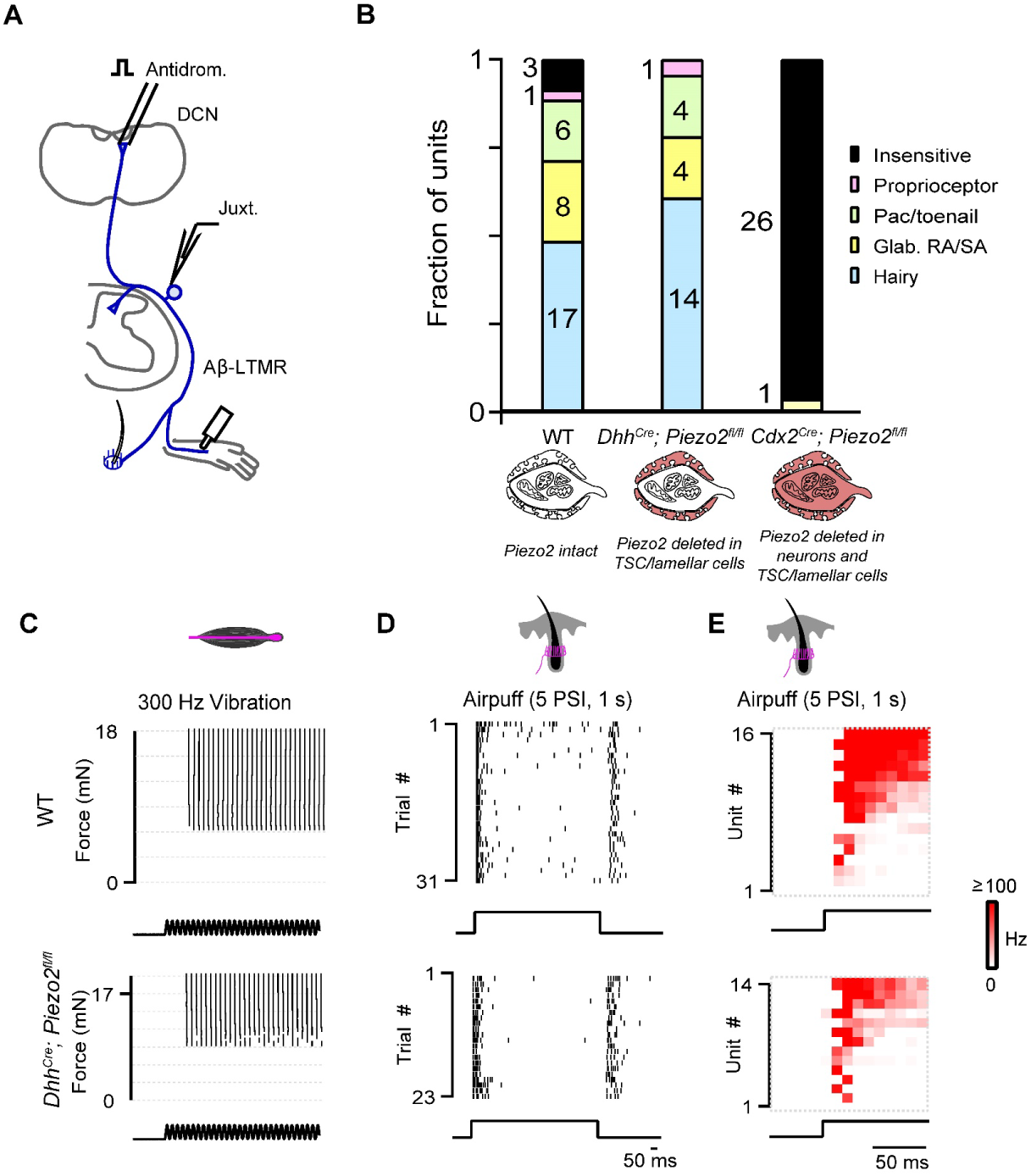
Deletion of Piezo2 in TSCs and lamellar cells does not affect light-touch responses in Aβ sensory neurons. (**A**) Schematic of juxtacellular recordings. Aβ sensory neurons were identified with antidromic activation of the dorsal column nucleus (DCN) or dorsal column (DC). Mechanical stimuli were applied to the hindpaw and hairy thigh to characterize the response properties of the units activated by DCN stimulation. (**B**) The fraction of DCN-projecting units identified as a specific sensory neuron class based on the location of their receptive field and response to low-threshold mechanical stimuli in wild type animals (n=3), *Dhh^Cre^;Piezo2^fl/fl^* mice (selective deletion of Piezo2 in peripheral Schwann cells, n=3), and *Cdx2^Cre^;Piezo2^fl/fl^*mice (deletion of Piezo2 in peripheral Schwann cells and sensory neurons, n=2). (**C**) Raster of single Pacinian Aβ RA-LTMR responses to vibration (300 Hz) delivered over a range of forces in a wild type animal (top) and one from a *Dhh^Cre^;Piezo2^fl/fl^* animal (bottom). (**D**) Raster of single lanceolate-forming Aβ-LTMR responses to air puff in a wild type animal (top) and a *Dhh^Cre^;Piezo2^fl/fl^* animal (bottom). (**E**) Histograms of airpuff responses for all identified Aβ hairy units activated by DCN stimulation in wild type animals (top) and *Dhh^Cre^;Piezo2^fl/fl^* animals (bottom). In the wild type animals, 15/16 hairy units were air puff-sensitive Aβ-LTMRs. In the *Dhh^Cre^;Piezo2^fl/fl^* animals, 14/15 hairy units were air puff-sensitive Aβ-LTMRs.

**Extended Data Figure 5.**
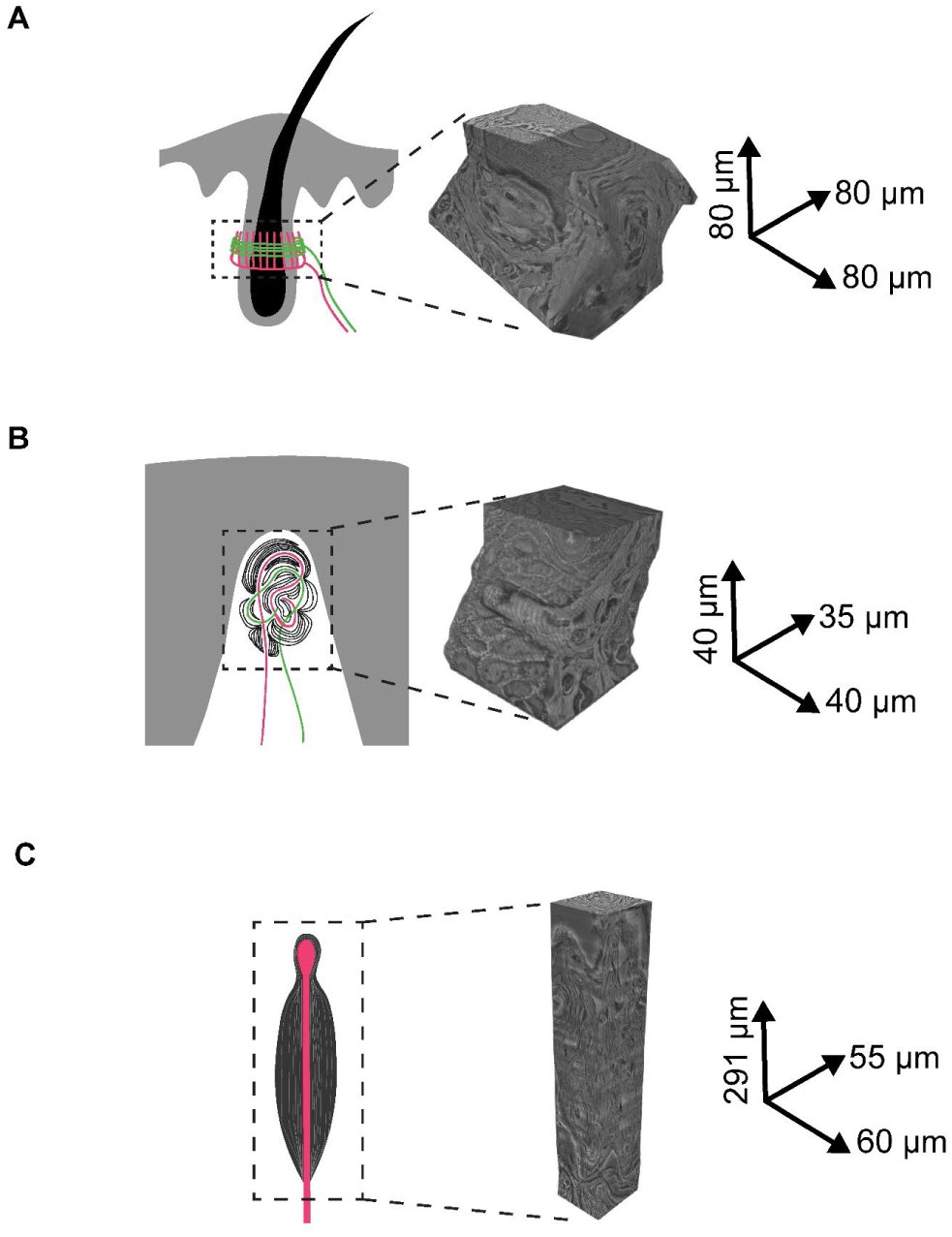
Aligned FIB-SEM volumes of a mouse guard hair, Meissner corpuscle, and Pacinian corpuscle. Full FIB-SEM volume with global alignment correction and dimensions for the (**A**) hairy skin sample containing a single guard hair; (**B**) glabrous skin sample containing a single Meissner corpuscle with two myelinated afferents; (**C**) sample containing a single Pacinian corpuscle.

**Extended Data Figure 6.**
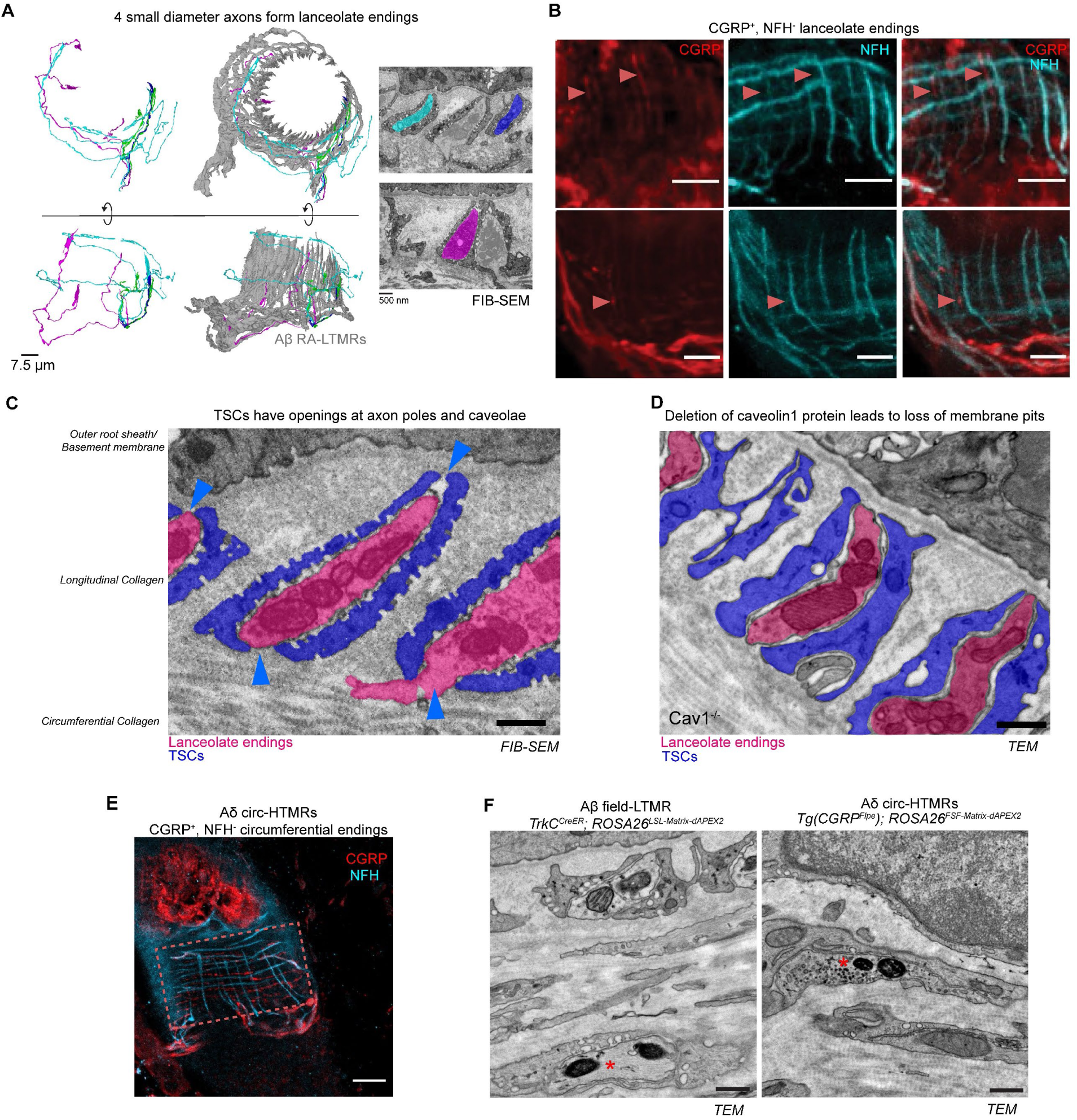
Characterization of cellular and structural features of the guard hair follicle. (**A**) 3D renderings of the six Aβ RA-LTMRs (gray) that form 47 lanceolate endings and four small caliber axons (magenta, green, cyan, and blue) that form six lanceolate endings around the guard hair. On right, single sections from the FIB-SEM volume with pseudo-colored lanceolate endings formed by Aβ RA-LTMRs and the small caliber neurons. (**B**) CGRP and NFH staining of hairy skin reveals CGRP^+^, NFH^-^ lanceolate endings, marked by red arrowheads. Examples from two animals are shown. Scalebar, 10 µm. (**C**) Single section from FIB-SEM volume with pseudo-colored Aβ RA-LTMR lanceolate ending (magenta) and TSCs (blue). The membrane of TSCs contains cup-shaped caveolae. Blue arrowheads indicate the gaps in TSC coverage that form closest to the outer root sheath and at the boundary of the longitudinal and circumferential collagen matrix. Scalebar, 500 nm. (**D**) Single section TEM of a hair follicle from a *Caveolin-1* knock-out animal with lanceolate endings pseudo-colored in magenta and TSCs pseudo colored in blue. This experiment was repeated in three animals. Scalebar, 500 nm. (**E**) CGRP and NFH staining of hairy skin showing CGRP^+^, NFH^-^ circumferential endings of Aδ circ-HTMRs. Scalebar, 20 µm. (**F**) Left, single section TEM showing mitochondrial dAPEX2 labeled Aβ field-LTMR (asterisk) from a *TrkC^CreER^; ROSA26^LSL-Matrix-dAPEX2^* animal treated with 0.5 mg of TAM at P5. Abundant neurofilaments were observed in these profiles. Right, single section TEM showing mitochondrial dAPEX2 labeled Aδ circ-HTMR (asterisk) from a *Tg(CGRP-Flpe); ROSA26^FSF-Matrix-dAPEX2^* animal. Neurofilaments were never found in these profiles, and these profiles tend to be smaller in diameter than Aβ field-LTMRs. Labeling was performed in one animal for each genotype. Scalebar, 500 nm.

**Extended Data Figure 7.**
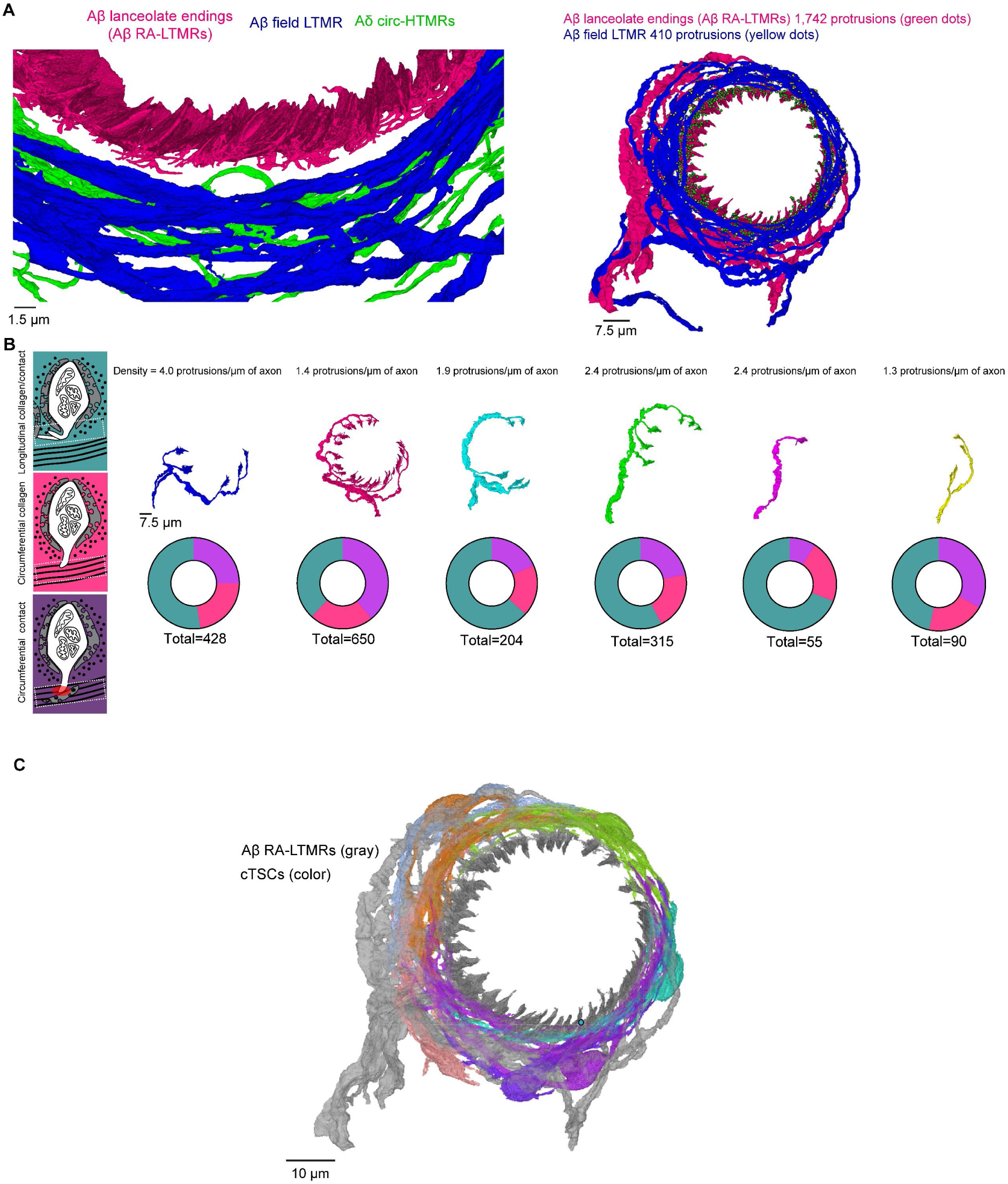
Axon protrusions are present along all Aβ RA-LTMR lanceolate endings and are minimally present along the less sensitive Aβ field LTMRs and Aδ circ-HTMRs. (**A**) Left, high magnification, 3D renderings from the guard hair FIB-SEM volume showing numerous axon protrusions along the Aβ RA-LTMR lanceolate endings (magenta) and the lack of protrusions along the Aβ field-LTMRs (blue) and Aδ circ-HTMRs (green). Right, 3D renderings showing all the protrusions along the Aβ RA-LTMR lanceolate endings (protrusions marked by green dots, Aβ RA-LTMRs rendered in magenta) and all the protrusions along the Aβ field-LTMRs (protrusions marked by yellow dots, Aβ field-LTMRs rendered in blue). (**B**) Quantification of axon protrusions for each of the six Aβ RA-LTMR axon segments. The density of protrusions for each axon segment is noted. Left schematic shows the three different termination patterns quantified. Teal: protrusions that stay within the local longitudinal collagen matrix. Magenta: protrusions that extend into the circumferential collagen matrix but do not contact cells in the circumferential collagen matrix. Purple: protrusions that extend into the circumferential collagen matrix and contact cells in the circumferential collagen matrix. A 3D render for each Aβ RA-LTMR is shown and the breakdown of protrusion terminals is shown below. (**C**) 3D rendering of Aβ RA-LTMRs in gray and circumferential TSCs (cTSCs) shown in color. A single teal dot marks the point of contact between an axon protrusion and a circumferential TSC.

**Extended Data Figure 8.**
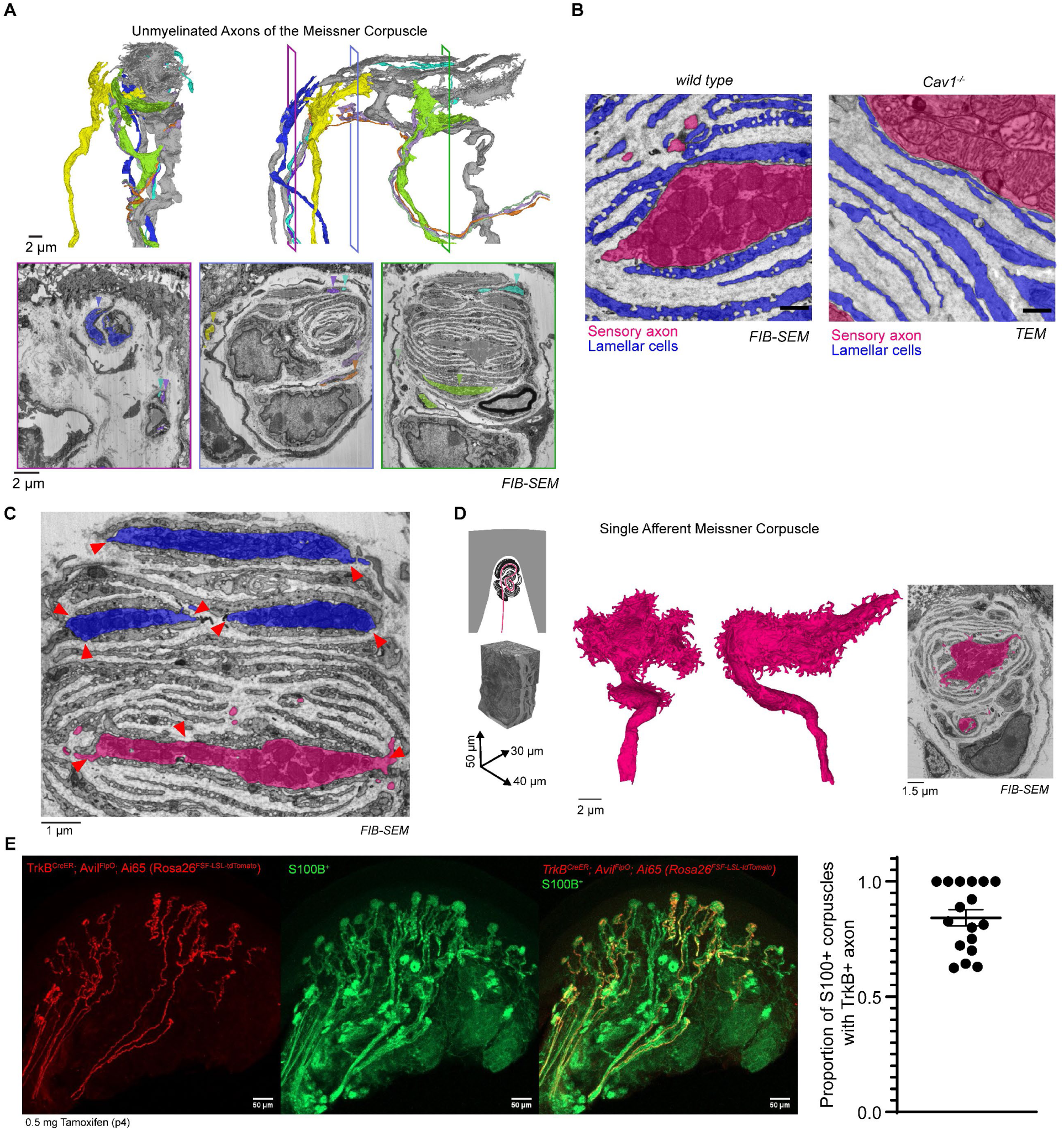
Characterization of cellular and structural features of the Meissner corpuscle. (**A**) 3D renderings of the two Aβ RA-LTMRs (shown in gray) and the seven (shown in color) small caliber/unmyelinated axons of the Meissner corpuscle. Three images taken from different depths of the Meissner corpuscle FIB-SEM volume show the pseudo-colored axon profiles of all seven small caliber/unmyelinated axons with arrowheads indicating their location. (**B**) Left, single section from FIB-SEM volume with Aβ RA-LTMR pseudo-colored magenta and lamellar cell processes pseudo-colored blue. The membranes of lamellar cells contain cup-shaped caveolae. Right, single section TEM image of a Meissner corpuscle in a *Caveolin-1* knock-out animal shows the loss of caveolae in the lamellar cell processes. This loss of caveolae was observed in the Meissner corpuscles of two *Caveolin-1* knock-out animals. Scalebar, 500 nm. (**C**) Single section from the FIB-SEM volume with the two Aβ RA-LTMRs shown in blue and magenta. Arrowheads indicate gaps in lamellar cell coverage of the sensory axons. (**D**) Left, schematic and FIB-SEM volume with global alignment correction and dimensions of a Meissner corpuscle innervated by a single Aβ RA-LTMR. Middle, 3D rendering of the Aβ RA-LTMR reconstructed from the Meissner corpuscle with a single myelinated afferent. Right, a single section from the FIB-SEM volume with the Aβ RA-LTMR pseudo-colored in magenta. (**E**) Co-staining thick (200 µm) sections from *TrkB^CreER^; Avil^FlpO^; Ai65 (Rosa26^FSF-LSL-tdTomato^)* animals with mCherry and S100B was used to quantify the proportion of S100B^+^ Meissner corpuscles innervated by a TrkB^+^ axon. Each data point represents the proportion of corpuscles with a TrkB^+^ axon in a single 200 µm thick section Mean ± SEM is plotted. Sections were collected from two animals.

**Extended Data Figure 9.**
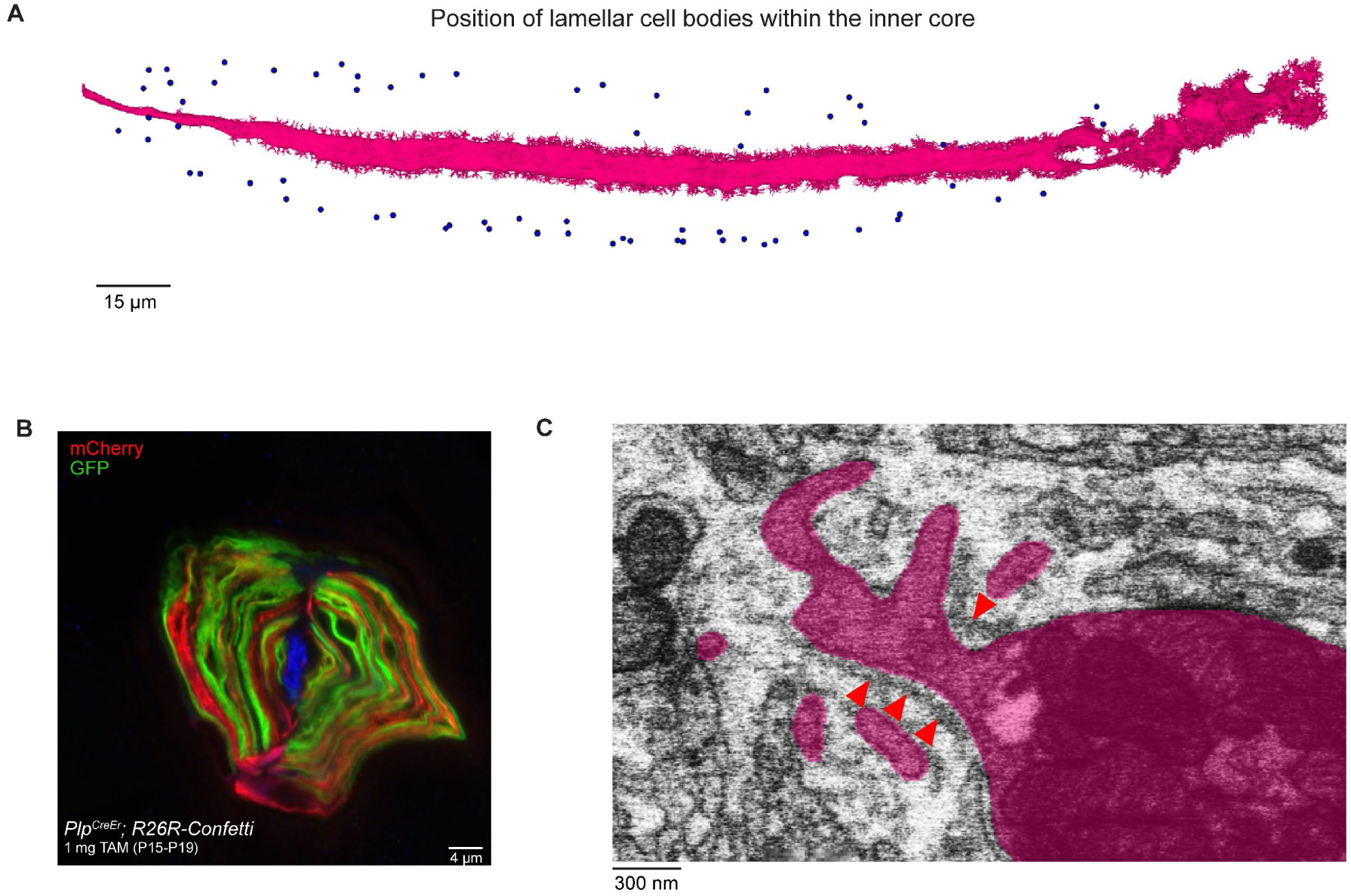
Characterization of cellular and structural features of the Pacinian corpuscle. (**A**) 3D rendering of the Aβ RA-LTMR innervating the Pacinian corpuscle. The blue dots mark the position of the soma of the lamellar cell that make up the inner core. (**B**) Cross section of a Pacinian corpuscle isolated from a *Plp^CreER^; R26R^Confetti^* mouse showing intermingled red and green processes from numerous lamellar cells. This experiment was performed in 1 animal. (**C**) Single section from the FIB-SEM volume of the Pacinian corpuscle with the Aβ RA-LTMR pseudo-colored magenta. Red arrowheads mark collagen fibers that surround the base of protrusions.

## REFERENCES

1. Handler, A. & Ginty, D. D. The mechanosensory neurons of touch and their mechanisms of activation. Nat. Rev. Neurosci. 22, 521–537 (2021).

2. Iggo, B. A. & Ogawa, H. Correlative physiological and morphological studies of rapidly adapting mechanoreceptors in cat’s glabrous skin. 266, 275–296 (1977).

3. Lynn, B. & Carpenter, S. E. Primary afferent units from the hairy skin of the rat hind limb. Brain Res. 238, 29–43 (1982).

4. Li, L. et al. The Functional Organization of Cutaneous Low-Threshold Mechanosensory Neurons. Cell 147, 1615–1627 (2011).

5. Lechner, S. G. & Lewin, G. R. Hairy sensation. Physiology 28, 142–150 (2013).

6. Koltzenburg, M., Stucky, C. L. & Lewin, G. R. Receptive properties of mouse sensory neurons innervating hairy skin. J. Neurophysiol. 78, 1841–1850 (1997).

7. Ranade, S. S. et al. Piezo2 is the major transducer of mechanical forces for touch sensation in mice. Nature 516, 121–125 (2014).

8. von Buchholtz, L. J. et al. Decoding Cellular Mechanisms for Mechanosensory Discrimination. Neuron 109, 285–298.e5 (2021).

9. Chesler, A. . et al. The role of PIEZO2 in human mechanosensation. N. Engl. J. Med. 375, 1355–1364 (2016).

10. Lehnert, B. P. et al. Mechanoreceptor synapses in the brainstem shape the central representation of touch. Cell 184, 5608–5621.e18 (2021).

11. Chirila, A. M. et al. Mechanoreceptor signal convergence and transformation in the dorsal horn flexibly shape a diversity of outputs to the brain. Cell 185, 4541–4559.e23 (2022).

12. Coste, B. et al. Piezo1 and Piezo2 are essential components of distinct mechanically activated cation channels. Science (80-. ). 330, 55–60 (2010).

13. Jin, P., Jan, L. Y. & Jan, Y. N. Mechanosensitive Ion Channels: Structural Features Relevant to Mechanotransduction Mechanisms. Annu. Rev. Neurosci. 43, 207–229 (2020).

14. Coste, B. et al. Piezo proteins are pore-forming subunits of mechanically activated channels. Nature 483, 176–181 (2012).

15. Kefauver, J. M., Ward, A. B. & Patapoutian, A. Discoveries in structure and physiology of mechanically activated ion channels. Nature 587, 567–576 (2020).

16. Loewenstein, W. & Mendelson, M. Components of receptor adaptation in a Pacinian corpuscle. J. Physiol. 177, 377–397 (1965).

17. Neubarth, N. L. et al. Meissner corpuscles and their spatially intermingled afferents underlie gentle touch perception. Science *(80-. ).* 368, 1–12 (2020).

18. Werner, G. & Mountcastle, V. B. Neural activity in mechanoreceptive cutaneous afferents: Stimulus-response relations, Weber functions, and information transmission. J. Neurophysiol. 28, 359–397 (1965).

19. Iggo, A. & Muir, A. The structure and function of a slowly adapting touch corpuscle in hairy skin. 1882, 763–796 (1969).

20. Woo, S. et al. Piezo2 is required for Merkel-cell mechanotransduction. Nature 509, 622– 626 (2014).

21. Woo, S.-H., Lumpkin, E. A. & Patapoutian, A. Merkel cells and neurons keep in touch. Trens Cell Biol 25, 74–81 (2015).

22. Ikeda, R. et al. Merkel cells transduce and encode tactile stimuli to drive aβ-Afferent impulses. Cell 157, 664–675 (2014).

23. Maksimovic, S. et al. Epidermal Merkel cells are mechanosensory cells that tune mammalian touch receptors. Nature 509, 617–621 (2014).

24. Nakatani, M., Maksimovic, S., Baba, Y. & Lumpkin, E. A. Mechanotransduction in epidermal Merkel cells. Pflugers Arch. Eur. J. Physiol. 467, 101–108 (2015).

25. Viswanathan, S. et al. High-performance probes for light and electron microscopy. Nat. Methods 12, 568–576 (2015).

26. Abdo, H. et al. Specialized cutaneous schwann cells initiate pain sensation. Science 365, 695–699 (2019).

27. Nikolaev, Y. A. et al. Lamellar cells in Pacinian and Meissner corpuscles are touch sensors. Sci. Adv. 6, (2020).

28. Schwaller, F. et al. USH2A is a Meissner’s corpuscle protein necessary for normal vibration sensing in mice and humans. Nat. Neurosci. 24, 74–78 (2020).

29. Andres, K. H. Über die Feinstruktur der Rezeptoren an Sinushaaren. Zeitschrift für Zellforsch. und Mikroskopische Anat. 75, 339–365 (1966).

30. Halata, Z. Sensory innervation of the hairy skin (light-and electronmicroscopic study). J. Invest. Dermatol. 101, 75S–81S (1993).

31. Takahashi-Iwanaga, H. Three-dimensional microanatomy of longitudinal lanceolate endings in rat vibrissae. J. Comp. Neurol. 426, 259–269 (2000).

32. Spencer, P. S. & Schaumburg, H. H. An ultrastructural study of the inner core of the Pacinian corpuscle. J. Neurocytol. 2, 217–35 (1973).

33. Takahashi-Iwanaga, H. The three-dimensional microanatomy of Meissner corpuscles in monkey palmar skin. J. Neurocytol. 32, 363–371 (2003).

34. García-Mesa, Y. et al. Merkel cells and Meissner’s corpuscles in human digital skin display Piezo2 immunoreactivity. J. Anat. 231, 978–989 (2017).

35. Pickles, J. O., Comis, S. D. & Osborne, M. P. Cross-links between stereocilia in the guinea pig organ of Corti, and their possible relation to sensory transduction. Hear. Res. 15, 103–112 (1984).

36. Howard, J. & Hudspeth, A. J. Compliance of the hair bundle associated with gating of mechanoelectrical transduction channels in the Bullfrog’s saccular hair cell. Neuron 1, 189–199 (1988).

37. Qiu, X. & Müller, U. Sensing sound: Cellular specializations and molecular force sensors. Neuron 110, 3667–3687 (2022).

38. Xu, C. S. et al. Enhanced FIB-SEM systems for large-volume 3D imaging. Elife 6, (2017).

39. Nguyen, T. M. et al. Structured cerebellar connectivity supports resilient pattern separation. Nature 613, 543–549 (2023).

40. Kuehn, E. D., Meltzer, S., Abraira, V. E., Ho, C. Y. & Ginty, D. D. Tiling and somatotopic alignment of mammalian low-threshold mechanoreceptors. Proc. Natl. Acad. Sci. U. S. A. 116, 9168–9177 (2019).

41. Li, L. & Ginty, D. D. The structure and organization of lanceolate mechanosensory complexes at mouse hair follicles. Elife 3, 1–24 (2014).

42. Parton, R. G. & Del Pozo, M. A. Caveolae as plasma membrane sensors, protectors and organizers. Nat. Rev. Mol. Cell Biol. 14, 98–112 (2013).

43. Bai, L. et al. Genetic Identification of an Expansive Mechanoreceptor Sensitive to Skin Stroking. Cell 163, 1783–1795 (2015).

44. Ghitani, N. et al. Specialized Mechanosensory Nociceptors Mediating Rapid Responses to Hair Pull. Neuron 95, 944–954.e4 (2017).

45. Sharma, N. et al. The emergence of transcriptional identity in somatosensory neurons. Nature 577, 392–398 (2020).

46. Zhang, Q., Lee, W. C. A., Paul, D. L. & Ginty, D. D. Multiplexed peroxidase-based electron microscopy labeling enables simultaneous visualization of multiple cell types. Nat. Neurosci. 22, 828–839 (2019).

47. Paré, M., Elde, R., Mazurkiewicz, J. E., Smith, A. M. & Rice, F. L. The Meissner corpuscle revised: A multiafferented mechanoreceptor with nociceptor immunochemical properties. J. Neurosci. 21, 7236–7246 (2001).

48. Zelena, J. Nerves and Mechanoreceptors. (Springer Netherlands, 1994).

49. Luo, W., Enomoto, H., Rice, F. L., Milbrandt, J. & Ginty, D. D. Molecular Identification of Rapidly Adapting Mechanoreceptors and Their Developmental Dependence on Ret Signaling. Neuron 64, 841–56 (2009).

50. Vega, J. A. et al. The inner-core, outer-core and capsule cells of the human pacinian corpuscles: An Immunohistochemical study. Eur. J. Morphol. 32, 11–8 (1994).

51. Talbot, W. H., Darian-Smith, I., Kornhuber, H. H. & Mountcastle, V. B. The sense of flutter-vibration: comparison of the human capacity with response patterns of mechanoreceptive afferents from the monkey hand. J. Neurophysiol. 31, 301–334 (1968).

52. Mountcastle, V. B., Talbot, W. H., Dar-Smith, I. & Kornhuber, H. H. Neural basis of the sense of flutter-vibration. Science *(80-. ).* 155, 597–600 (1967).

53. Kaidoh, T. & Inoué, T. Intercellular junctions between palisade nerve endings and outer root sheath cells of rat vellus hairs. J. Comp. Neurol. 420, 419–427 (2000).

54. Kaidoh, T. & Inoué, T. N-Cadherin Expression in Palisade Nerve Endings of Rat Vellus Hairs. J. Comp. Neurol. 506, 525–534 (2008).

55. Yap, A. S., Duszyc, K. & Viasnoff, V. Mechanosensing and mechanotransduction at cell – cell junctions. Cold Spring Harb. Perspect. Biol. 10, 1–16 (2018).

56. Mallon, B. S., Elizabeth Shick, H., Kidd, G. J. & Macklin, W. B. Proteolipid promoter activity distinguishes two populations of NG2-positive cells throughout neonatal cortical development. J. Neurosci. (2002). doi:10.1523/jneurosci.22-03-00876.2002

57. Doerflinger, N. H., Macklin, W. B. & Popko, B. Inducible site-specific recombination in myelinating cells. Genes. (United States) (2003). doi:10.1002/gene.10154

58. Choi, S. et al. Parallel ascending spinal pathways for affective touch and pain. Nature 587, 258–263 (2020).

59. Razani, B. et al. Caveolin-1 Null Mice Are Viable but Show Evidence of Hyperproliferative and Vascular Abnormalities. J. Biol. Chem. 276, 38121–38138 (2001).

60. Rutlin, M. et al. The cellular and molecular basis of direction selectivity of Aδ-LTMRs. Cell 159, 1640–1651 (2014).

61. Madisen, L. et al. A robust and high-throughput Cre reporting and characterization system for the whole mouse brain. Nat. Neurosci. 13, 133–140 (2010).

62. Coutaud, B. & Pilon, N. Characterization of a novel transgenic mouse line expressing Cre recombinase under the control of the Cdx2 neural specific enhancer. Genesis (2013). doi:10.1002/dvg.22421

63. Jaegle, M. et al. The POU proteins Brn-2 and Oct-6 share important functions in Schwann cell development. Genes Dev. 17, 1380–1391 (2003).

64. Huang, S., O’Donovan, K. J., Turner, E. E., Zhong, J. & Ginty, D. D. Extrinsic and intrinsic signals converge on the Runx1/CBFβ transcription factor for nonpeptidergic nociceptor maturation. Elife 4, 1–24 (2015).

65. Pang, S. & Xu, C. S. Methods of enhanced FIB-SEM sample preparation and image acquisition. in Methods in Cell Biology: Volume Electron Microscopy Volume 181 (ed. P Verkade)

66. Xu, C. S., Pang, S., Hayworth, K. J. & Hess, H. F. Transforming FIB-SEM Systems for Large-Volume Connectomics and Cell Biology. in Neuromethods (2020). doi:10.1007/978-1-0716-0691-9_12

67. Xu, C. S. et al. An open-access volume electron microscopy atlas of whole cells and tissues. Nature (2021). doi:10.1038/s41586-021-03992-4

68. Klein, S., Staring, M., Murphy, K., Viergever, M. A. & Pluim, J. P. W. Elastix: A toolbox for intensity-based medical image registration. IEEE Trans. Med. Imaging (2010). doi:10.1109/TMI.2009.2035616

69. Shamonin, D. P. et al. Fast parallel image registration on CPU and GPU for diagnostic classification of Alzheimer’s disease. Front. Neuroinform. (2014). doi:10.3389/fninf.2013.00050

70. Phelps, J. S. et al. Reconstruction of motor control circuits in adult Drosophila using automated transmission electron microscopy. Cell (2021). doi:10.1016/j.cell.2020.12.013

71. Kuan, A. T. et al. Dense neuronal reconstruction through X-ray holographic nano-tomography. Nat. Neurosci. (2020). doi:10.1038/s41593-020-0704-9

72. Funke, J. et al. Large Scale Image Segmentation with Structured Loss Based Deep Learning for Connectome Reconstruction. IEEE Trans. Pattern Anal. Mach. Intell. (2019). doi:10.1109/TPAMI.2018.2835450

73. Boergens, K. M. et al. WebKnossos: Efficient online 3D data annotation for connectomics. Nat. Methods (2017). doi:10.1038/nmeth.4331

74. Funke, J. gunpowder: A library to facilitate machine learning on multi-dimensional images. (2020). Available at: https://github.com/funkey/gunpowder/.

75. Kisuk Lee, Jonathan Zung, Peter Li, Viren Jain, H. S. S. Superhuman accuracy on the SNEMI3D connectomics challenge. arXiv Prepr. (2017).

76. Nguyen, T., Malin-Mayor, C., Patton, W. & Funke, J. Daisy: block-wise task dependencies for luigi. (2022).

