## Supplementary Table of Contents for "Three-dimensional reconstructions of mechanosensory end organs suggest a unifying mechanism underlying dynamic, light touch"

**Supplementary Video 1: Reconstructed Aβ rapidly adapting low-threshold mechanoreceptors (RA-LTMRs) that form lanceolate endings around a mouse guard hair.**
3D volumetric reconstructions of the six Aβ RA-LTMR axon segments that form the 47 lanceolate endings that surround the mouse guard hair.

**Supplementary Video 2: Reconstruction of Aβ RA-LTMRs and a single terminal Schwann cell (TSC) around a mouse guard hair.**
3D volumetric reconstructions of the Aβ RA-LTMR sensory neurons (magenta) and a single TSC (blue).

**Supplementary Video 3: Reconstruction of Aβ RA-LTMRs and all associated terminal Schwann cells (TSCs) around a mouse guard hair.**
3D volumetric reconstructions of the Aβ RA-LTMR sensory neurons (white) and all TSCs that associate with them to form the lanceolate endings that surround the mouse guard hair.

**Supplementary Video 4: Reconstruction of Aβ field-LTMRs and Aβ RA-LTMRs around a mouse guard hair.**
3D volumetric reconstructions of the two Aβ field-LTMR axons (blue and green) within the circumferential collagen matrix that surround the Aβ RA-LTMR sensory neurons (white) that lie adjacent to the hair shaft.

**Supplementary Video 5: Reconstruction of Aδ circumferential high-threshold mechanoreceptors (circ-HTMRs) and Aβ RA-LTMRs around a mouse guard hair.**
3D volumetric reconstructions of the two Aδ circ-HTMR axons (purple and green) within the circumferential collagen matrix that surround the Aβ RA-LTMR sensory neurons (white) that lie adjacent to the hair shaft.

**Supplementary Video 6: Reconstruction of a subset of circumferential support cells (CSCs) and Aβ RA-LTMRs around a mouse guard hair.**
3D volumetric reconstructions of four CSCs (green, blue, purple, and cyan) that reside within the circumferential collagen matrix that surround the Aβ RA-LTMR sensory neurons (white) that lie adjacent to the hair shaft.

**Supplementary Video 7: Axon protrusions from Aβ RA-LTMR lanceolate endings frequently contact CSCs.**
Raw FIB-SEM images from a portion of the mouse guard hair volume showing axon protrusions from Aβ RA-LTMR sensory neurons (magenta) extending into the circumferential collagen matrix and contacting CSCs (green). TSCs are shown in blue.

**Supplementary Video 8: Two reconstructed Aβ sensory axons innervating a Meissner corpuscle from mouse glabrous skin.**
3D volumetric reconstructions of two Aβ sensory axons that innervate a single Meissner corpuscle isolated from the forepaw digit tip of a mouse.

**Supplementary Video 9: Axon protrusions extend from the Aβ RA-LTMR of the Meissner corpuscle and frequently contact lamellar cell processes.**
Raw FIB-SEM images from a portion of the mouse Meissner corpuscle volume showing axon protrusions from the putative $TrkB^{+}$ Aβ RA-LTMR (magenta) that extend into the local collagen matrix and frequently contact nearby lamellar cells (yellow, cyan, and green).

**Supplementary Video 10: Reconstructed Aβ RA-LTMR of the Pacinian corpuscle.**
3D volumetric reconstruction of the Aβ RA-LTMR that innervates a Pacinian corpuscle isolated from the periosteum surrounding the fibula of a mouse's hindleg.

**Supplementary Video 11: Axon protrusions extend from the Aβ RA-LTMR of the Pacinian corpuscle and frequently contact lamellar cell processes.**
Raw FIB-SEM images from a portion of the ultraterminal region of the Pacinian corpuscle volume showing axon protrusions that extend into the local collagen matrix and frequently contact nearby lamellar cells.
